# Computing coalescence rates for complex demographies and sampling configurations

**DOI:** 10.64898/2026.04.09.717519

**Authors:** Jiatong Liang, Jonathan Terhorst

## Abstract

Inference of population history from genetic data relies, implicitly or explicitly, on the distribution of coalescence times, because population size changes, migration, and admixture all leave characteristic signatures in genealogies. The distribution of pair-wise coalescent rates in particular has emerged as a popular target for demographic inference methods. However, pairwise coalescent rates have limited power to resolve recent history, because recent coalescences in samples of size two are rare. In this article, we introduce `demestats`, a software library for computing first-coalescence and cross-coalescence rate functions for structured demographic models specified in the `demes` format. The method computes the instantaneous rate at which the first coalescence event occurs conditional on no prior coalescence for arbitrary sampling configurations, combines exact calculations with mean-field approximations for larger samples, and is differentiable with respect to model parameters. In simulations, these statistics recover recent population size change and recent migration more accurately than pairwise summaries. Applied to tree sequences inferred from the 1000 Genomes Project, the method estimates recent growth rates and present-day effective population sizes across five human superpopulations.

## 1 Introduction

Large whole-genome datasets (Auton et al., 2015; Sudlow et al., 2015) and recent progress in ancestral recombination graph (ARG) inference (Deng et al., 2025; Kelleher et al., 2019a; Speidel et al., 2019) have made it increasingly practical to analyze reconstructed genealogies directly (Lewanski et al., 2024). Historically, coalescent-based demographic inference methods have treated underlying coalescence times as latent and attempted to integrate them out, either explicitly or through summary statistics such as the site frequency spectrum (SFS) (Dilber and Terhorst, 2024; Kamm et al., 2020, 2017; Li and Durbin, 2011; Schiffels and Durbin, 2014; Terhorst et al., 2017). Reconstructed tree sequences now provide estimated genealogies whose node times can serve as noisy proxies, or approximate posterior samples, for latent coalescence times. Demographic inference can therefore use these times directly rather than summaries of present-day variation.

A basic object in this setting is the instantaneous coalescent rate (ICR), which is the rate at which sampled lineages coalesce at time *t* in the past, conditioned on no coalescence at a more recent time. In a panmictic population, the reciprocal of the ICR is proportional to the effective population size *N*_*e*_ (Waples, 2022). Under population structure, this simple interpretation breaks down: the ICR depends not only on demographic parameters, but also on how the lineages were sampled. Inference from ICR curves therefore amounts to fitting the observed distribution of coalescence times by the family of distributions induced by a demographic model.

Several earlier works have developed this idea. Mazet et al. (2016) introduced the pairwise inverse instantaneous coalescent rate (IICR) and showed that, under population structure, the distribution of pairwise coalescence times can always be re-expressed as an apparent history of population size change under panmixia. Chikhi et al. (2018) gave a simulation-based construction for the pairwise IICR by simulating many independent *T*_2_ values and estimating the corresponding survival and hazard functions empirically. Rodríguez et al. (2018) extended this to the non-stationary structured coalescent, giving a transition matrix method for exact pairwise ICR curves under time-varying migration and deme size changes. Arredondo et al. (2021) and DeHaas et al. (2025) used pairwise ICR curves for demographic inference, and Chikhi et al. (2024) generalized the construction to samples of size *k >* 2 through the time to the first coalescent event, denoted ICR_*k*_. Related work has also used cross-coalescence events, meaning the first coalescent event between samples from different populations, to study recent population separation (Schiffels and Durbin, 2014). Most recently, Jouniaux et al. (2026) extended exact pairwise ICR computation to a much broader class of non-stationary structured models.

A main obstacle is that exact ICR calculations are still largely model specific. In practice, one typically has to derive a new ICR formula, or a new system of equations, for each new demographic model. The recent work of Jouniaux et al. (2026) addresses much of this problem for the pairwise case by giving a general construction for *k* = 2 under a broad class of non-stationary structured models. Extending this idea to larger samples is harder. For *k >* 2 and for arbitrary sampling configurations, the calculation must track many more possible arrangements of ancestral lineages among demes, and the underlying state of the system becomes a multidimensional array.

In this paper we introduce `demestats`, a Python library for computing first-coalescence and cross-coalescence statistics for demographic models specified in the `demes` format (Gower et al., 2022). The calculation follows sampled lineages backwards through the demographic history. Between population splits, pulses, and admixture events, it updates the distribution of their locations under migration and coalescence. At each event, it redistributes the lineages according to the event’s ancestry proportions. Parts of the demographic history that cannot yet interact are calculated separately and combined when migration or shared ancestry connects them. The structure of this algorithm is adapted from an algorithm used by the `momi` family of methods for computing the expected SFS (Dilber and Terhorst, 2024; Kamm et al., 2020, 2017). For large samples, exact computations become intractable, and we develop a mean-field approximation that tracks expected lineage counts rather than the full distribution. Our implementation is differentiable with respect to demographic parameters, which permits likelihood-based inference and other statistical calculations.

## 2 Methods

A demographic history divides backward time into intervals separated by events such as population splits, pulses, and changes in migration. Within an interval, the set of demes is fixed, although population sizes and migration rates may vary with time. Our algorithm alternates between following migration and coalescence within each time interval, and updating lineage locations at the event that marks the end of that interval. Lineages that cannot yet coalesce because of population structure can be treated independently. We call a set of demes whose lineages must be followed jointly a *block*. Each sampled deme begins in a separate block, and two blocks are combined when migration or shared ancestry first allows their lineages to enter the same deme. Until then, the joint distribution of their locations factors over the blocks. Their survival probabilities multiply, so their log survival probabilities add. For example, lineages sampled from two isolated descendant demes can be followed separately until the population split, when both sets of lineages enter their common ancestral deme.

An *event tree* records the blocks and the order in which they are joined. We adapt the event tree calculus developed for the `momi` series of SFS methods (Dilber and Terhorst, 2024; Kamm et al., 2020, 2017) to track lineage locations and coalescent probabilities in structured populations. Operations on the event tree move lineages between blocks or combine blocks. A `Merge` forms a joint state from two blocks that were previously independent. Moving backwards through a population split places lineages from a descendant deme into its ancestor; `Split1` and `Split2` perform this update when the affected demes are already in one block or still in two blocks, respectively. `Pulse` and `Admix` assign lineages to ancestral demes using the specified ancestry proportions. For additional information on these verbs and what they represent, see Kamm et al. (2020).

### 2.1 Instantaneous Coalescent Rate

For a sample of *k* lineages, let *T*_*k*_ denote the time to the first coalescent event and let 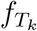 denote its density. Define

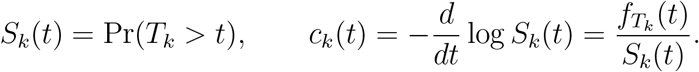

The instantaneous coalescent rate is ICR_*k*_(*t*) = *c*_*k*_(*t*). Its reciprocal is the IICR (Mazet et al., 2016).

Under panmixia, 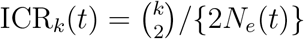, so it is in one-to-one correspondence with *N*_*e*_. Under structured models this correspondence no longer holds because the rate also depends on the ancestral locations of the lineages. Previous work derived exact calculations for increasingly broad classes of structured models (Chikhi et al., 2024; Jouniaux et al., 2026; Rodríguez et al., 2018).

Conditional on *T*_*k*_ *> t*, all *k* lineages are still present. Our algorithm works by tracking their ancestral locations. It uses one of two representations depending on the problem size. One representation records the count *x*_*i*_ in each deme *i*:

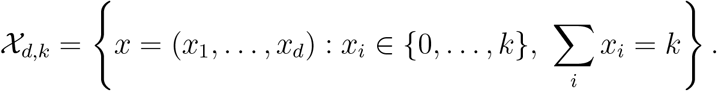

We refer to this as the count-by-deme or *occupancy* representation. A second representation records the current deme of each of the *k* labeled lineages, giving a state *z* = (*z*_1_, …, *z*_*k*_) *∈ {*1, …, *d}* ^*k*^. These two arrays have (*k* + 1)^*d*^ and *d*^*k*^ entries, respectively. A heuristic selects the smaller representation once for each exact calculation.

Demographic events act to alter distributions on these state spaces. Moving backwards across a population split places lineages from the descendant deme in its ancestor. At a pulse or admixture event, each affected lineage is assigned to an ancestral deme with the specified probability, so the number of lineages taking one ancestry is binomially distributed. If an event connects two blocks that were previously independent, their distributions are multiplied before this update. Appendix A defines these updates for both representations.

Between demographic events, let *X*(*t*) be the count-by-deme vector and define

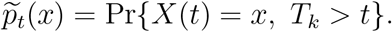

This quantity includes both the distribution of lineage locations and the probability that no coalescence has yet occurred. In particular,

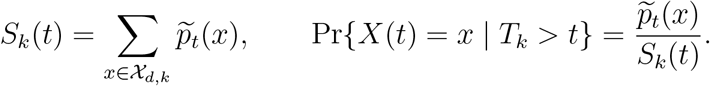

Let *M*_*ij*_(*t*) be the backward-time migration rate from deme *i* to deme *j* and *N*_*i*_(*t*) be the effective population size of deme *i* at time *t*. From state *x*, migration changes the state to *x e*_*i*_ +*e*_*j*_ at rate *x*_*i*_*M*_*ij*_(*t*), where *e*_*i*_ has one in coordinate *i* and zero elsewhere. Let *Q*_mig_(*t*) be the transition-rate matrix formed from these moves. Also write *η*_*i*_(*t*) = 1*/* 2*N*_*i*_(*t*) . In state *x*, the total rate of a first coalescence is

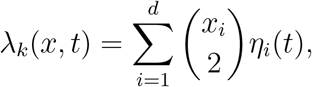

because any pair in the same deme can coalesce. If 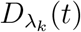 is the diagonal matrix with entries *λ*_*k*_(*x, t*), then the row vector 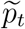 satisfies

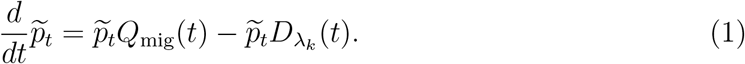

The first term moves probability among possible lineage locations. The second removes probability at the rate of the first coalescence. Averaging the rate over the configurations that have survived to time *t* gives

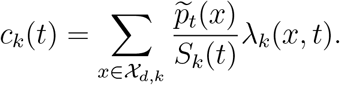

The algorithm returns this rate together with log *S*_*k*_(*t*).

Equation (1) has a simple solution on an interval *I* = [*t*_0_, *t*_1_) with no migration:

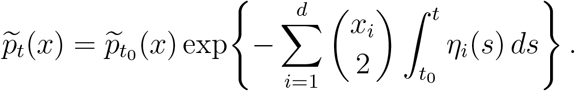

If migration is active but population sizes are constant on the interval, `demestats` uses the matrix exponential of 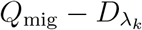 . If migration rates or population sizes vary with time, it solves Equation (1) numerically.

### 2.2 Cross Coalescence

Cross-coalescence rates (CCR) summarize coalescences between two groups of sampled lineages and can reveal recent population separation (Schiffels and Durbin, 2014). As with the ICR, our algorithm allows to compute these rates for arbitrary demographies and sampling configurations.

In a CCR computation, there are two demes, say red and blue. Let *T*_*×*_ be the time of the first coalescence between the demes. Define

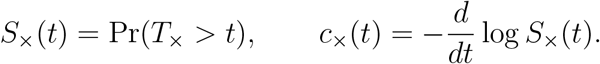

The CCR is *c*_*×*_(*t*), the rate of a red-blue coalescence conditional on no such event at a more recent time.

Let *R*_*i*_(*t*) and *B*_*i*_(*t*) be the numbers of red and blue lineages in deme *i* at time t. Coalescences between lineages of the same color may occur before *T*_*×*_, so these counts can decrease with time. For count vectors *ρ* and *β*, define

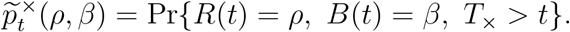

Summing this quantity over all count vectors gives *S*_*×*_(*t*). In a configuration (*ρ, β*), there are *ρ*_*i*_*β*_*i*_ possible red-blue pairs in deme *i*, so the rate of cross-coalescence is

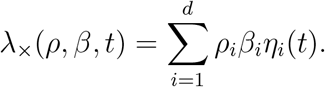

Between demographic events, 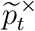 satisfies

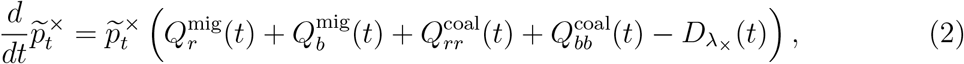

where the first two terms move red and blue lineages among demes, the next two terms move probability when two lineages of the same color coalesce, and 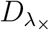 removes probability when the first red-blue coalescence occurs. Normalizing the survival probabilities and averaging the rate gives

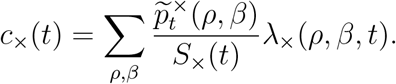

Appendix B gives the entries of the four transition-rate matrices in Equation (2).

At population splits, pulses, and admixture events, the rules for lineage locations described for ICR are applied separately to the two colors. When a `Merge` joins two previously independent blocks, their configuration probabilities are multiplied and their log-survival values are added.

### 2.3 Mean-field approximations

The exact calculations described above track the probability distribution of every possible lineage configuration. For large numbers of lineages, this exact calculation becomes computationally infeasible. To enable scaleable computations, we additionally developed a mean-field (MF) approximation that instead tracks some moments of this distribution.

One option could be to simply track the expected number of lineages in each deme at each time point, in place of the full joint distribution of their locations. However, it is easy to see that the resulting approximation would be very inaccurate, because coalescent rates depend on second order (pairwise) interactions. Consider one lineage sampled from each of two symmetrically migrating demes, *A* and *B*. Under symmetric migration without coalescence, the expected counts are (1, 1) for all time. Initially each deme contains exactly one lineage, so neither contains a pair. After the lineages have mixed, each deme contains zero, one, or two lineages with probabilities 1*/*4, 1*/*2, and 1*/*4, respectively. The expected counts are still (1, 1), but the expected number of pairs is now 1*/*4 in each deme. Substituting the expected count into *x*(*x −* 1)*/*2 would give zero both initially and after mixing, missing this change in the coalescence rate.

The missing information that accounts for the change in coalescence rate is the variance of the lineage counts. Write *T* for *T*_*k*_ in ICR and *T*_*×*_ in CCR, and similarly, write S for *S*_*k*_ and *S*_*×*_. If *L*_*p*_(*t*) is the lineage count in deme *p* and 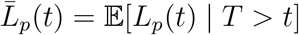, then

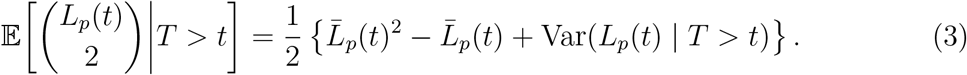

In the example above, the variance increases from zero to 1*/*2 in each deme, supplying the missing 1*/*4 expected number of pairs. For CCR, if *R*_*p*_(*t*) and *B*_*p*_(*t*) are the red and blue counts and 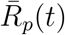 and 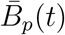 are their conditional means, the corresponding identity is

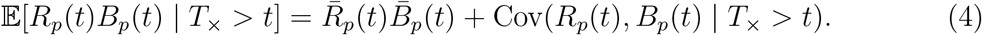

Thus both rates require second moments as well as expected counts.

To approximate these conditional moments, partition the sample into groups *g*, each containing *n*_*g*_ lineages sampled from the same deme and, for CCR, of the same color. Let *L*_*gp*_(*t*) be the random count from group *g* in deme *p*. We distinguish its exact conditional mean from its mean-field approximation by writing

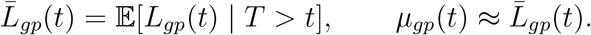

Thus *L*_*gp*_ denotes a random count in the exact process, 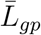 its exact conditional mean, and *µ*_*gp*_ the approximant that we compute.

Directly evolving conditional variances and covariances would require equations for still higher moments in both ICR and CCR, leading to a moment closure problem. We close the system by approximating the states of distinct lineages as independent conditional on survival. Within each group, the counts across demes are then multinomial, with deme probabilities *µ*_*gp*_(*t*)*/n*_*g*_; for CCR, an additional category accounts for lineages lost through same-color coalescences. In particular,

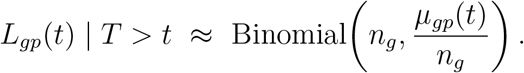

Summing the means and variances of these independent group counts gives

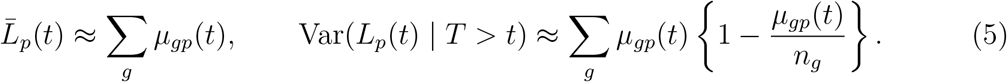

For CCR, Equation (5) applies separately to each color. Note that the red-blue covariance in Equation (4) is zero under the independence approximation, since the two counts sum over disjoint sets of lineages.

Substituting these moments into Equations (3) and (4) gives the approximate coalescent rates

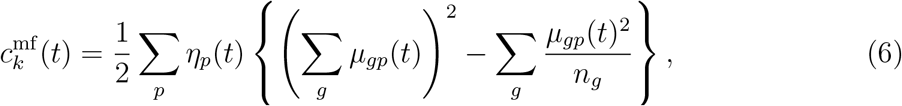

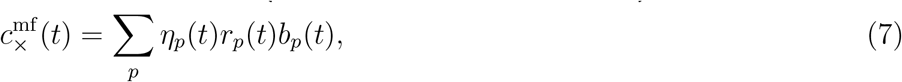

where *r*_*p*_(*t*) and *b*_*p*_(*t*) sum the approximate conditional means *µ*_*gp*_(*t*) over red and blue sampling groups, respectively.

#### 2.3.1 Time evolution of *µ*_*gp*_(*t*)

Now we describe how the approximate conditional means are evolved through time. We first consider ICR, with *T* = *T*_*k*_ and *S*(*t*) = Pr(*T*_*k*_ *> t*); the modifications necessary for the CCR case are described afterwards.

Fix a genealogy that has survived to time *t* and consider a short interval *dt*. A migration from deme *s* ≠ *p* into *p* occurs at rate *L*_*gs*_*M*_*sp*_ and increases the *L*_*gp*_ by one. The diagonal entries of *M*, whose rows sum to zero, account for departures. Thus migration contributes an expected change ∑*s L*_*gs*_*M*_*sp*_ *dt*. If the genealogy’s overall rate of coalescence is *λ*, its first coalescence occurs with probability *λ dt* + *o*(*dt*). This makes its contribution to the weighted count drop from *L*_*gp*_ to zero, giving an expected change *− L*_*gp*_*λ dt*. Suppressing time arguments, the expected change in the weighted count from this configuration is therefore

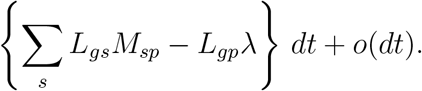

The survival indicator itself drops from one to zero at coalescence, so its expected change in a surviving configuration is *− λ dt* + *o*(*dt*). For a genealogy that has already coalesced, both the weighted count and the survival indicator remain zero. Taking expectations over the current configuration therefore gives the following changes over an infinitesimal interval:

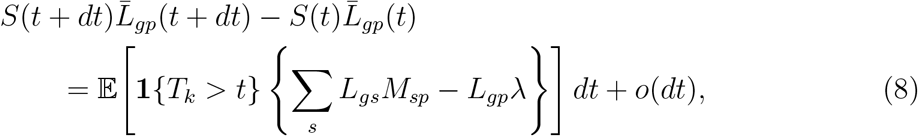

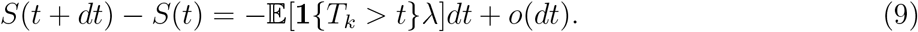

Now since

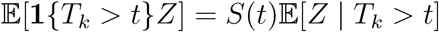

for any integrable random variable *Z*, we have

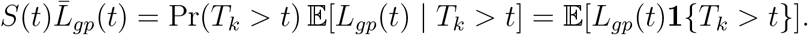

Plugging this in, dividing by *dt* and taking *dt →* 0 in (8) and (9) gives

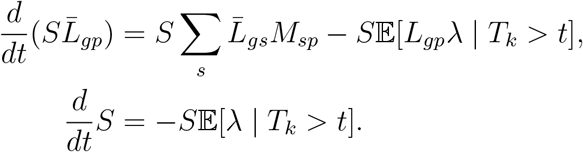

Applying the product rule to 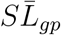 and dividing by *S* gives

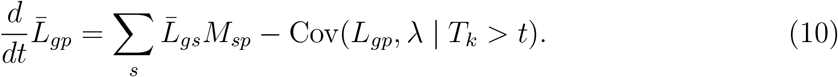

The covariance term subtracts the contribution of configurations that coalesce faster than average. For intuition, imagine two lineages with no migration: one is known to be in deme *A*, while the other could be in *A* or *B*. They can coalesce only if both are in *A*, so the longer they go without coalescing, the more likely the second lineage is to be in *B*.

To evaluate the covariance under the independence approximation, let *h*_*gp*_ be the expected rate at which a lineage from group *g* in deme *p* coalesces with another lineage. Writing *µ*_*p*_ = ∑*g µ*_*gp*_, we have

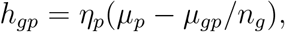

where the subtraction *µ*_*gp*_*/n*_*g*_ excludes pairing a lineage with itself. Now fix a lineage *l* from group *g*, whose probability of being in deme *p* is *q*_*gp*_ = *µ*_*gp*_*/n*_*g*_ under the approximation. Let *d*_*g*_ be the expected rate of coalescences that do not involve *l*. Independence makes this rate the same regardless of the location of *l*. The expected total rate is therefore *d*_*g*_ + *h*_*gp*_ given that *l* is in *p*. Averaged over possible locations for *l* this becomes 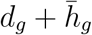, where 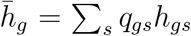. Each group member contributes *q*_*gp*_(*d*_*g*_ + *h*_*gp*_) to E[*L*_*gp*_*λ T*_*k*_ *> t*], and 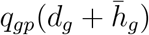 to 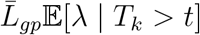. Summing their differences over the *n*_*g*_ group members gives

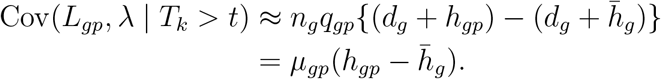

Replacing 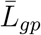 by *µ*_*gp*_ in (10) and using this covariance approximation, we get

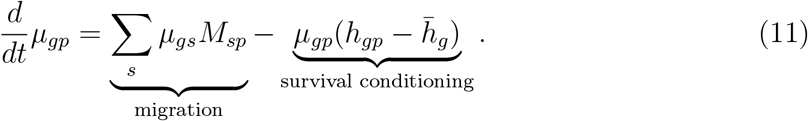

For CCR, replace *T*_*k*_ by *T*_*×*_ and let *λ* and *d*_*g*_ include only red–blue coalescences. Then *h*_*gp*_ = *η*_*p*_*b*_*p*_ for a red group, or *h*_*gp*_ = *η*_*p*_*r*_*p*_ for a blue group. The covariance calculation is otherwise the same. Same-color coalescences can also occur before *T*_*×*_, removing lineages from histories that still satisfy the survival condition. At such a coalescence, the ancestor inherits either descendant’s sampling group with probability 1*/*2. Under the independence approximation, a red lineage from group *g* has *r*_*p*_ *− µ*_*gp*_*/n*_*g*_ possible red partners on average. This contributes an additional loss term on the right hand side of Equation (11) in the amount of

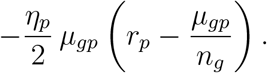

For a blue group, replace *r*_*p*_ by *b*_*p*_ in this loss term.

#### 2.3.2 Demographic events

At a demographic event, the approximate conditional means in each row of *µ*(*t*) are updated using the specified ancestry probabilities. Here, *t*^*−*^ := lim_*ε↓*0_(*t − ε*) and *t*^+^ := lim_*ε↓*0_(*t* + *ε*). If *A*_*pa*_ is the probability that a lineage in deme *p* enters ancestral deme *a*, then

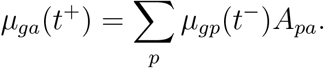

A `Merge` joins two independent blocks by placing their count matrices on the diagonal of a larger matrix and adding their log-survival values:

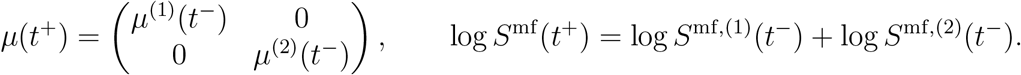

For a `Pulse` of proportion *α* from source deme *s* into destination deme *d*, backward-time ancestry gives

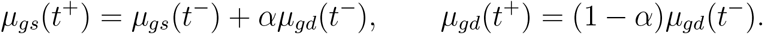

The remaining demes and log survival are unchanged. A `Split1` adds the descendant column to its ancestral column; `Split2` first joins the two blocks and then performs this addition. An `Admix` divides the child column among its ancestors using their ancestry proportions.

### 2.4 Local identifiability analysis

Our implementation supports automatic differentiation. Once the first-coalescence density

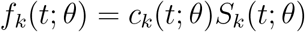

is available, we can differentiate it with respect to the demographic parameters *θ*. Writing

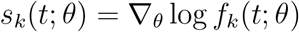

for the score contributed by one observation at time *t*, the Fisher information for a single first-coalescence time is

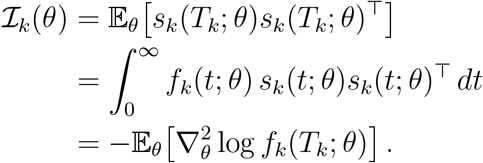

For *n* independent draws from the same *T*_*k*_ distribution, the total information is *n ℐ*_*k*_(*θ*).

Suppose a parameter *θ*_*m*_ first affects the demographic history at time *a*_*m*_. Observations with *t < a*_*m*_ carry no information about that parameter. Observations at later times may remain informative even after the interval or event in which the parameter appears. The diagonal entry of the Fisher Information matrix for this parameter is

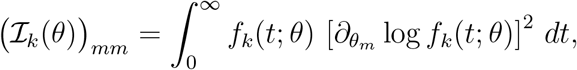

which vanishes whenever either *f*_*k*_(*t*; *θ*) = 0 or 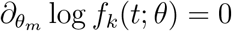. The former means that no first-coalescence mass is present at time *t*, while the latter means that the density is insensitive to *θ*_*m*_.

We evaluate ℐ_*k*_(*θ*) at a specified demography, approximate the matrix integral by trapezoidal quadrature on a time grid, and use automatic differentiation to obtain the needed derivatives of the exact ICR curve (Bradbury et al., 2018; Press et al., 2007). Diagonal entries measure sensitivity to individual parameters, and off-diagonal entries describe local dependence between parameter effects. For each figure, we divide all plotted diagonal entries by the largest one, so the largest displayed value is one. These values are relative sensitivities for the stated parameterization and should not be interpreted as an absolute scale of identifiability. We associate each value with its corresponding entry in the original `demes` model and display the result with a modified version of `demesdraw` (Gower, 2023).

## 3 Results

### 3.1 Local parameter sensitivity in example demographies

The Fisher information measures how strongly the distribution of the observed data changes with a model parameter near a specified demography. We summarize this local sensitivity using the normalized diagonal entries defined in Section 2.4 and plot them on the demographic graph. A score of one marks the most sensitive parameter among those shown in a figure; a score near zero means that the chosen first-coalescence distribution changes very little with that parameter. We first apply this diagnostic to several `stdpopsim` models (Lauterbur et al., 2023).

In the single-deme model with exponential growth `Africa_1T12`, for *k* = 2, sensitivity is greatest in the constant epoch older than 5,920 generations and negligible in the growth epoch within the last 204.6 generations (Figure 1). With *k* = 20, the strongest sensitivity shifts to the intermediate epoch (204.6–5,920 generations ago), the recent growth epoch becomes more informative, and the oldest epoch becomes weak. In the single-deme `Zigzag_1S14` model, sensitivity at *k* = 2 declines steadily toward the present. At *k* = 20, it instead concentrates in the two recent zigzag epochs spanning 133–533 and 33.3–133 generations ago, while the two oldest epochs have weak local sensitivity.

**Figure 1:**
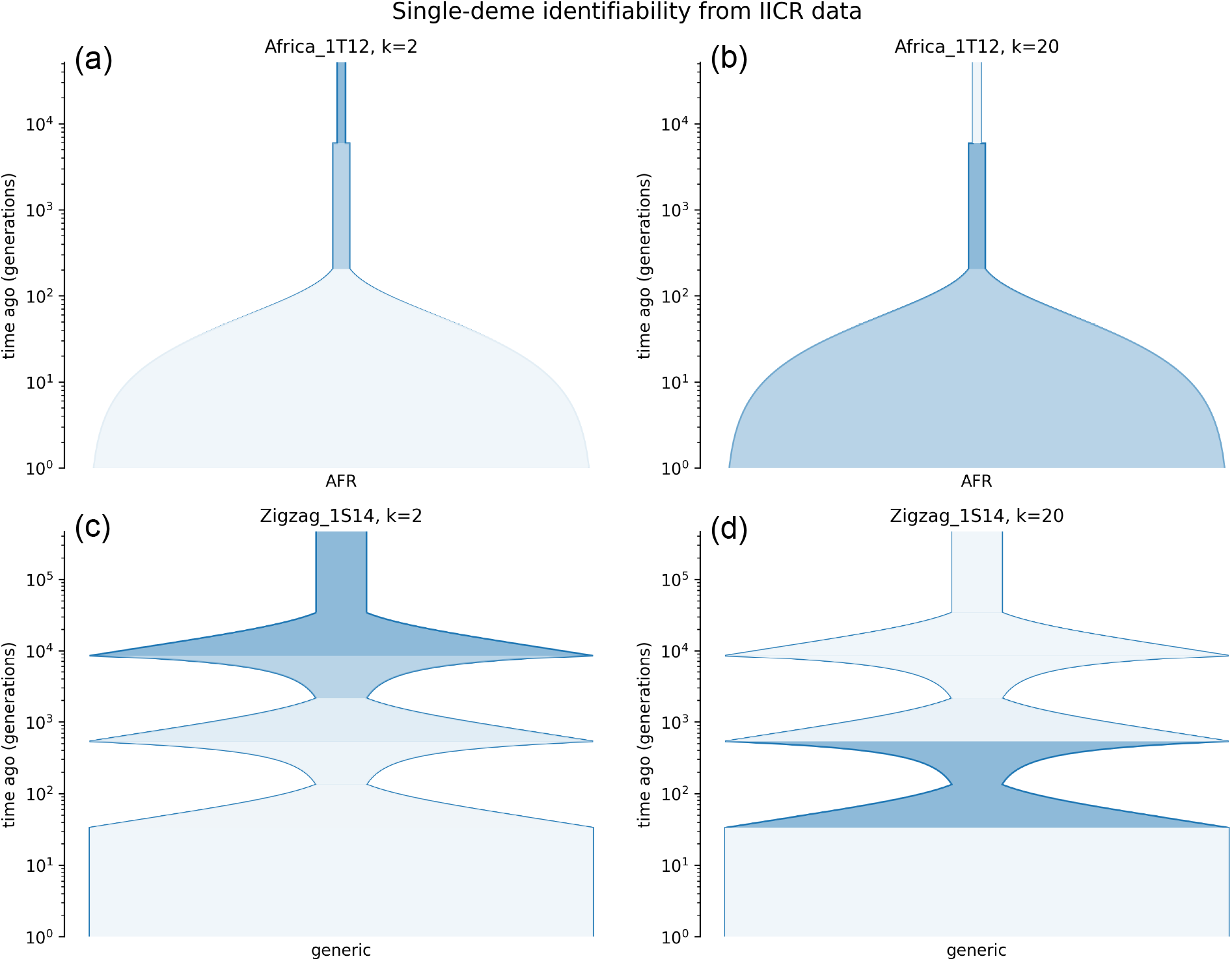
Relative local sensitivity for two single-deme `stdpopsim` models. Colored tube regions correspond to size epochs with relatively high Fisher information, while white regions indicate weak local sensitivity. In both `Africa_1T12` (a)-(b) and `Zigzag_1S14` (c)-(d), increasing *k* shifts the strongest sensitivity from ancient epochs toward more recent size changes.

To test whether the pairwise sensitivity plot predicts the behavior of an actual *k* = 2 inference procedure, we simulated 10 independent 100 Mb diploid chromosomes under `Zigzag_1S14` and ran the pairwise sequentially Markovian coalescent (PSMC) method with the standard human settings of Li and Durbin (Li and Durbin, 2011). The left panel of Figure 2 repeats the pairwise sensitivity plot for `Zigzag_1S14`, while the right panel shows the 10 inferred size histories with the same epochwise shading mapped onto time. The sensitivity plot predicts a sharp limit for pairwise data in this model. Only the two oldest epochs, older than 8533 generations, carry non-negligible information according to our method: the ancestral constant epoch has normalized score 1.0 and the second-oldest expansion has score 0.21, whereas all more recent epochs fall below 0.09. The PSMC reconstructions follow this pattern closely: across replicates, they consistently recover a broad ancient excursion around 10^4^ generations and the older ancestral plateau, but they do not reproduce the three recent sawtooth oscillations between 33 and 8533 generations. Quantitatively, the median absolute log_10_ error of the inferred histories is 0.053 in the informative ancient band versus 0.259 in the recent band, corresponding to typical multiplicative errors of 1.13x and 1.82x, respectively.

**Figure 2:**
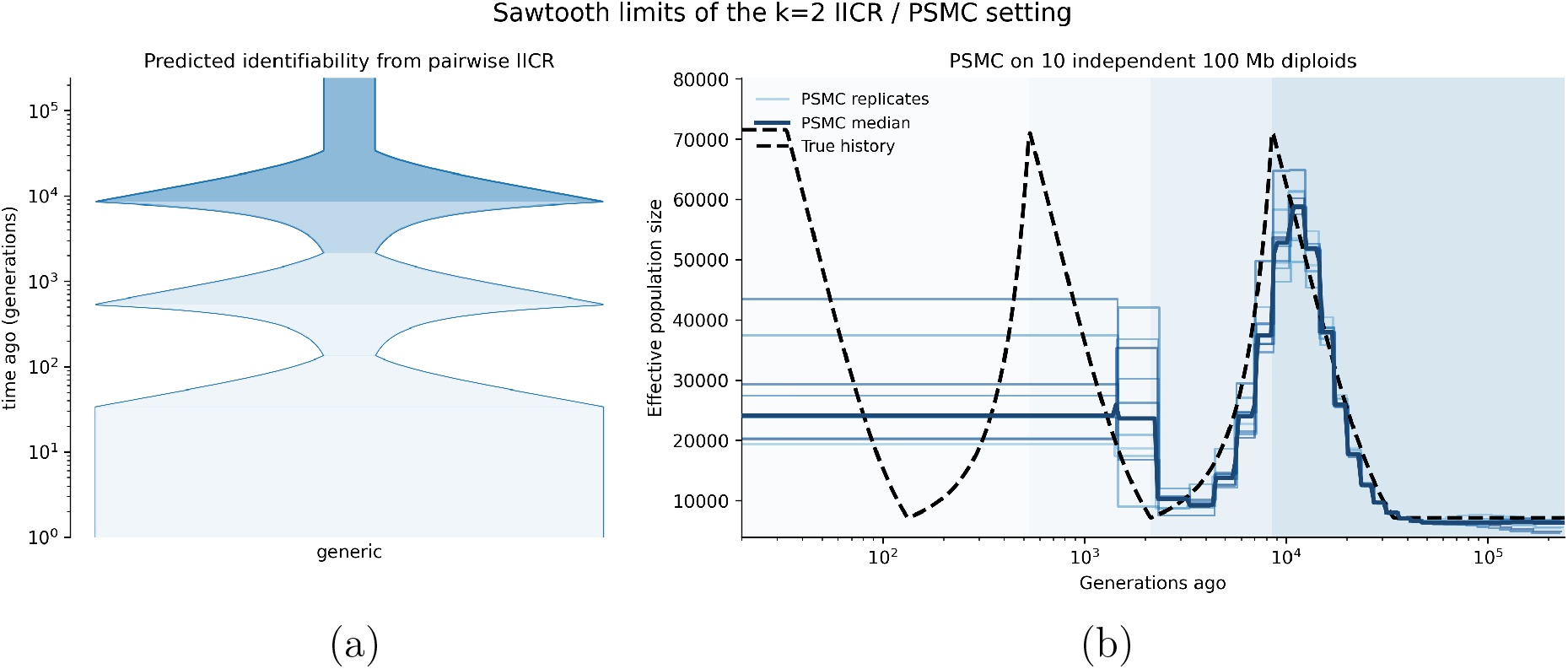
Pairwise limitations in the `stdpopsim Zigzag_1S14` sawtooth model. (a) The *k* = 2 identifiability sensitivity plot obtained from the diagonal of the ICR Fisher information matrix. (b) PSMC reconstructions from 10 independent 100 Mb simulated diploid chromosomes under the same model. The dashed black curve is the true size history, and the background shading on the right reuses the same epochwise Fisher scores as the left panel. PSMC tracks the informative ancient band but smooths over the recent oscillations assigned low sensitivity by the pairwise calculation.

#### 3.1.1 A cross-population sample in OutOfAfrica_3G09

We used `OutOfAfrica_3G09` for a cross-population example. We chose a sample that contained one lineage from the Utah residents with European ancestry (CEU) and one from the Han Chinese in Beijing (CHB). There is little opportunity for coalescence inside the modern descendant demes, so the distribution has little local sensitivity to their sizes. Figure 3 shows that all scores for descendant size epochs are numerically negligible, with the largest normalized size score below 4 *×* 10^*−*14^. Instead, the largest diagonal entries of the Fisher information matrix correspond to migration rates, especially the older migration from Yoruba in Ibadan (YRI) to CEU between 848 and 5,600 generations ago, followed by the reciprocal `CEU` *→* `YRI` rate on the same interval and the recent `CHB` *→*`CEU` rate within the last 848 generations. The largest sensitivities therefore correspond to migration pathways that can bring the two lineages into the same deme; the left panel highlights the three most sensitive migration intervals.

**Figure 3:**
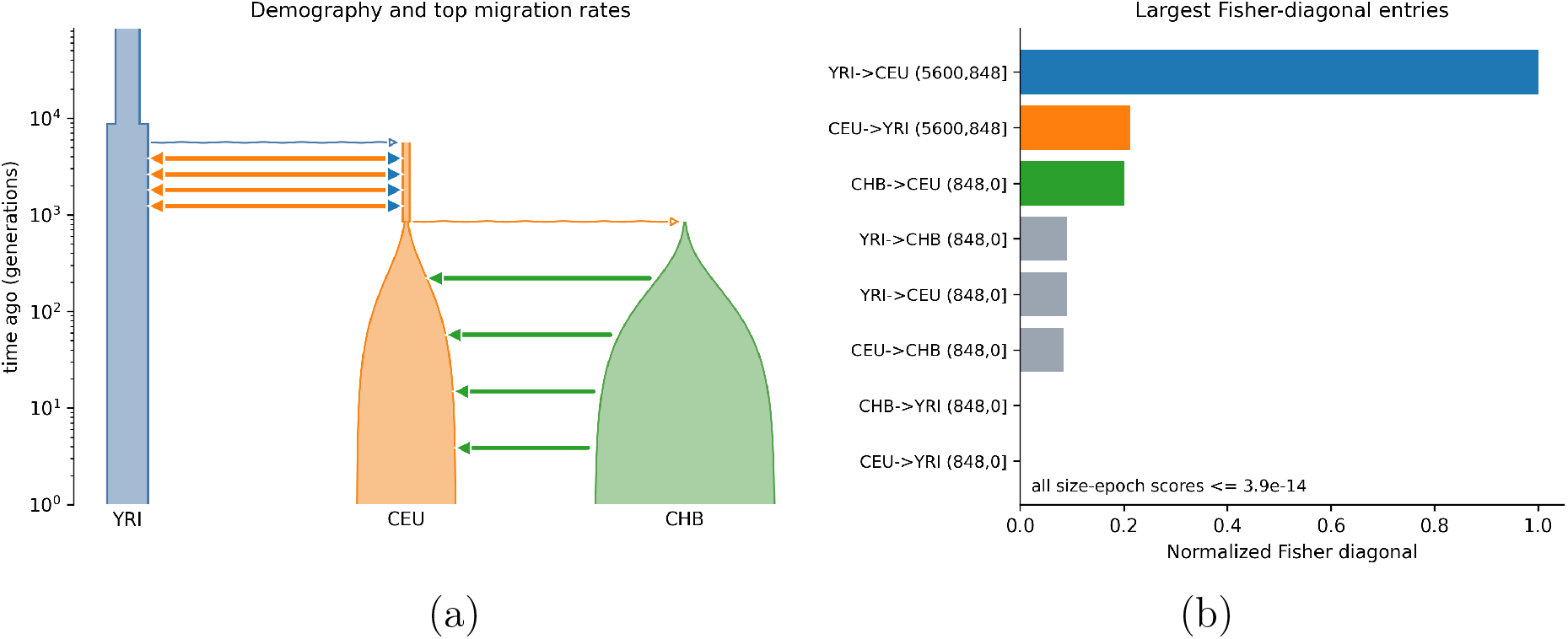
Relative local sensitivity in the `stdpopsim OutOfAfrica_3G09` model for one CEU and one CHB lineage. (a) Demographic graph with the three migration rates having the largest diagonal entries of the Fisher information matrix highlighted; the corresponding bars on the right use the same colors. (b) The largest normalized Fisher diagonal entries. For this cross-population configuration, the largest sensitivities are assigned to migration rates; all scores for descendant size epochs are below 4 *×* 10^*−*14^.

### 3.2 Mean-field accuracy and sampling configuration

We compared the mean-field (MF) approximation described in Section 2.3 with the exact solver. We evaluated both hazards on the same time grid where the exact calculation remained feasible and summarized the discrepancy by

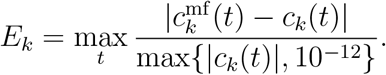

In a symmetric two-deme migration model with balanced sampling, correlations among lineage locations are weak and the mean-field approximation is already tight at small sample sizes. Figure 4a shows near overlap between the exact and mean-field hazards for *k* = 2, 10, and 25, with max relative errors of 1.24%, 0.95%, and 0.79%, respectively. In this exchangeable setting, increasing *k* improves the approximation because the lineage counts in the two demes remain well mixed and first moments capture most of the relevant variation.

**Figure 4:**
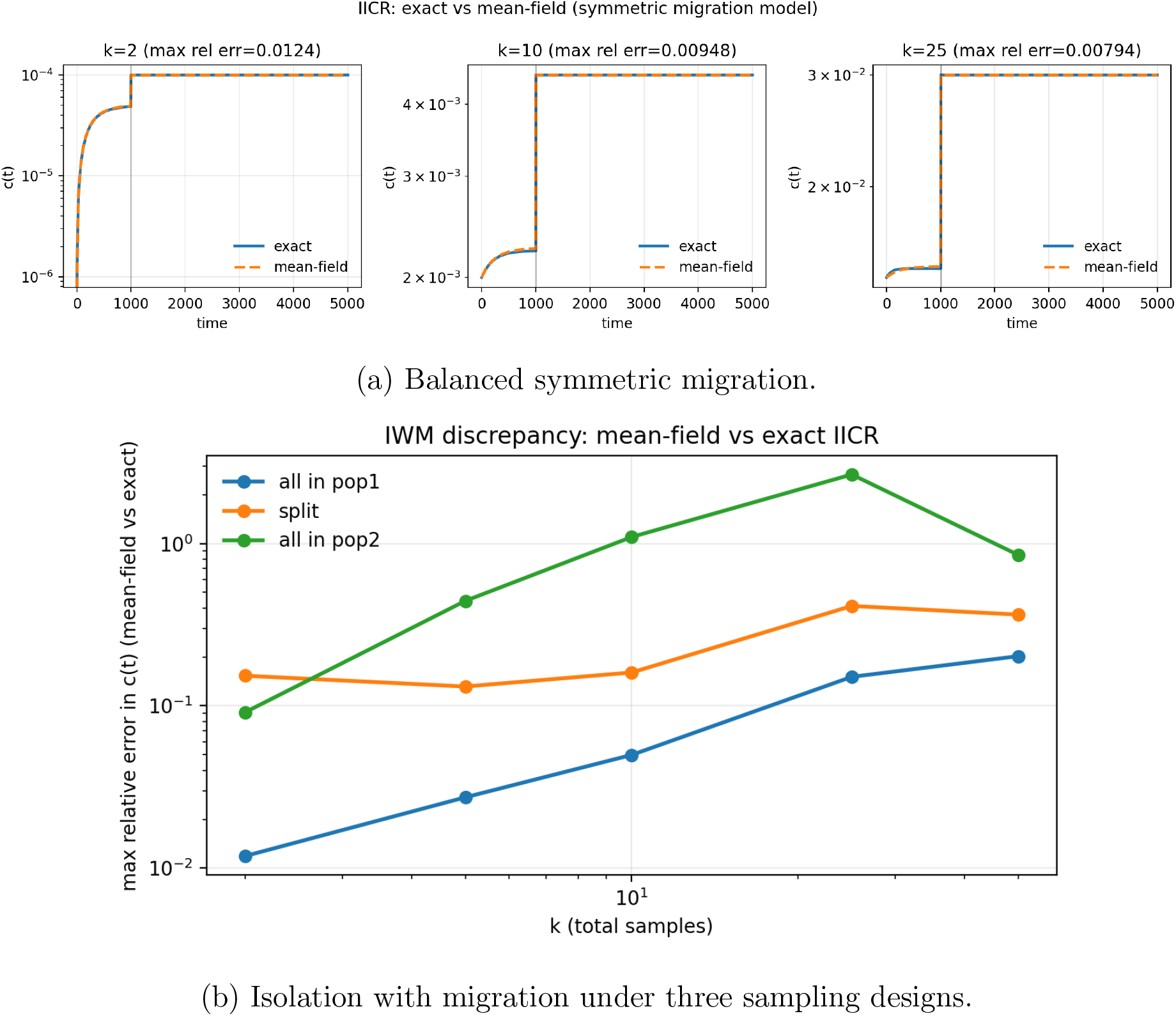
(a) Exact versus mean-field ICR comparisons. In a symmetric two-deme migration model with balanced sampling between demes, the mean-field hazard closely tracks the exact hazard and improves with *k*. (b) In the IwM model, the max relative hazard error *E*_*k*_ depends strongly on where the lineages are sampled: errors remain moderate for balanced samples, whereas concentrating all samples in the smaller descendant population produces much larger discrepancies.

The isolation-with-migration (IwM) model reveals a different regime in which error can accumulate depending on the sampling configuration. We evaluated an example IwM model with descendant population sizes 4000 and 1000, split time 1000 generations, and asymmetric migration rates 10^*−*4^ and 2 *×* 10^*−*4^. Figure 4b shows that when all lineages are sampled from the larger descendant population (`pop1`), the discrepancy remains modest, rising from 1.19% at *k* = 2 to 20.2% at *k* = 50. For an even split between descendant populations, the error stays moderate, between 13.2% and 41.3%. However, when all lineages are concentrated in the smaller descendant population (`pop2`), the discrepancy becomes much larger: 44.3% at *k* = 5, 110% at *k* = 10, and 266% at *k* = 25. The approximation can therefore improve with *k* in balanced settings yet deteriorate when many lineages are forced through a small deme, because rapid within-deme coalescences result in correlations that are dropped by our algorithm (Section 2.3). Mean-field is therefore safest for diffuse or balanced sampling designs, whereas highly concentrated sampling in a small deme should use the exact solver whenever the state space is still tractable.

### 3.3 Recent migration and sample size

We quantified how larger samples improve inference of very recent migration in an IwM model. We fixed ancestral size 10,000, descendant sizes 15,000 and 20,000, split time *T*_*s*_ = 30 generations, and symmetric recent migration rate *m*. We then sampled *n* red lineages from population 1 and *n* blue lineages from population 2, so *k* = 2*n*. Under the null model *H*_0_ : *m* = 0, a red-blue coalescence before the split is impossible. Under the alternative *H*_1_ : *m >* 0, any event with *T*_*×*_ *< T*_*s*_ must have been created by recent migration. Hence, the probability that one marginal tree contains an informative cross-coalescence is

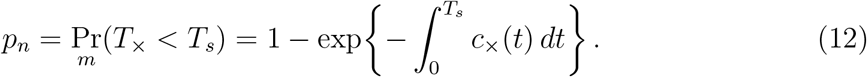

For *L* independent marginal trees, the most powerful test against *H*_0_ rejects whenever at least one tree has *T*_*×*_ *< T*_*s*_,

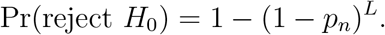

Thus, by studying how *p*_*n*_ scales with sample size, we can quantify power to detect very recent migration.

For small *m* and times *t < T*_*s*_, the total migration flux is proportional to *nm*, because each of the *n* lineages in a population can migrate independently. Once a migrant arrives, it sees *O*(*n*) lineages of the opposite color as potential cross-coalescence partners. Consequently, the cross-coalescence hazard at early times satisfies

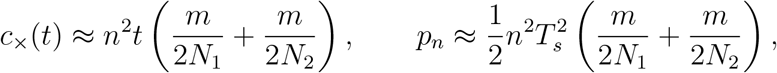

where the approximation for *p*_*n*_ also uses 1 *− e*^*−x*^ *≈ x* for small *x*. Thus, the gain from larger samples grows approximately quadratically in the number of lineages per population. Figure 5 confirms this scaling: before the population split, the probability of cross-coalescence increases rapidly with *n*. For the small-*n* cases where the exact calculation is tractable, the mean-field approximation matches *p*_*n*_ in Equation (12) to within a few percent.

**Figure 5:**
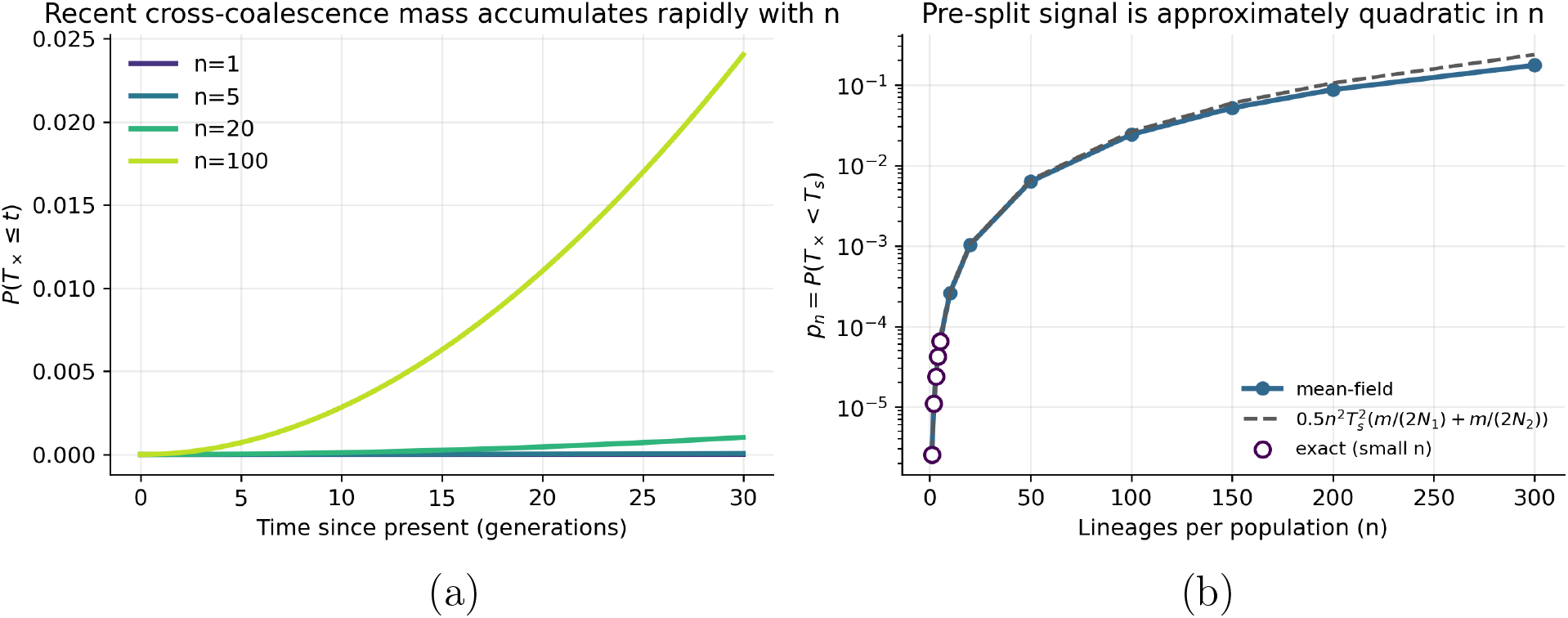
First coalescence probability in IwM model with *T*_*s*_ = 30 generations and symmetric migration rate *m* = 10^*−*4^. (a) The cumulative probability *P* (*T*_*×*_ *≤ t*) for balanced samples of size *n* per population. Larger samples substantially increase the probability of cross-coalescence before the population split. (b) The probability that the first cross-coalescence occurs before the split as a function of *n*. The dashed curve is given by the approximate formula. Open circles show exact CCR values for the small-*n* cases where the exact computation is tractable.

For *m* = 10^*−*4^, the probability increases from 2.6 *×* 10^*−*6^ at *n* = 1 to 0.174 at *n* = 300. The number of independent trees required for 80% power therefore collapses from 6.2 *×* 10^5^ at *n* = 1 to 2.5 *×* 10^4^ at *n* = 5, 255 at *n* = 50, 66 at *n* = 100, and about 9 at *n* = 300. Figure 6 shows the same effect as full power curves. For a threefold weaker migration rate *m* = 3 *×* 10^*−*5^, increasing the sample from *n* = 5 to *n* = 300 reduces the 80% threshold from roughly 8.2 *×* 10^4^ trees to 28.

**Figure 6:**
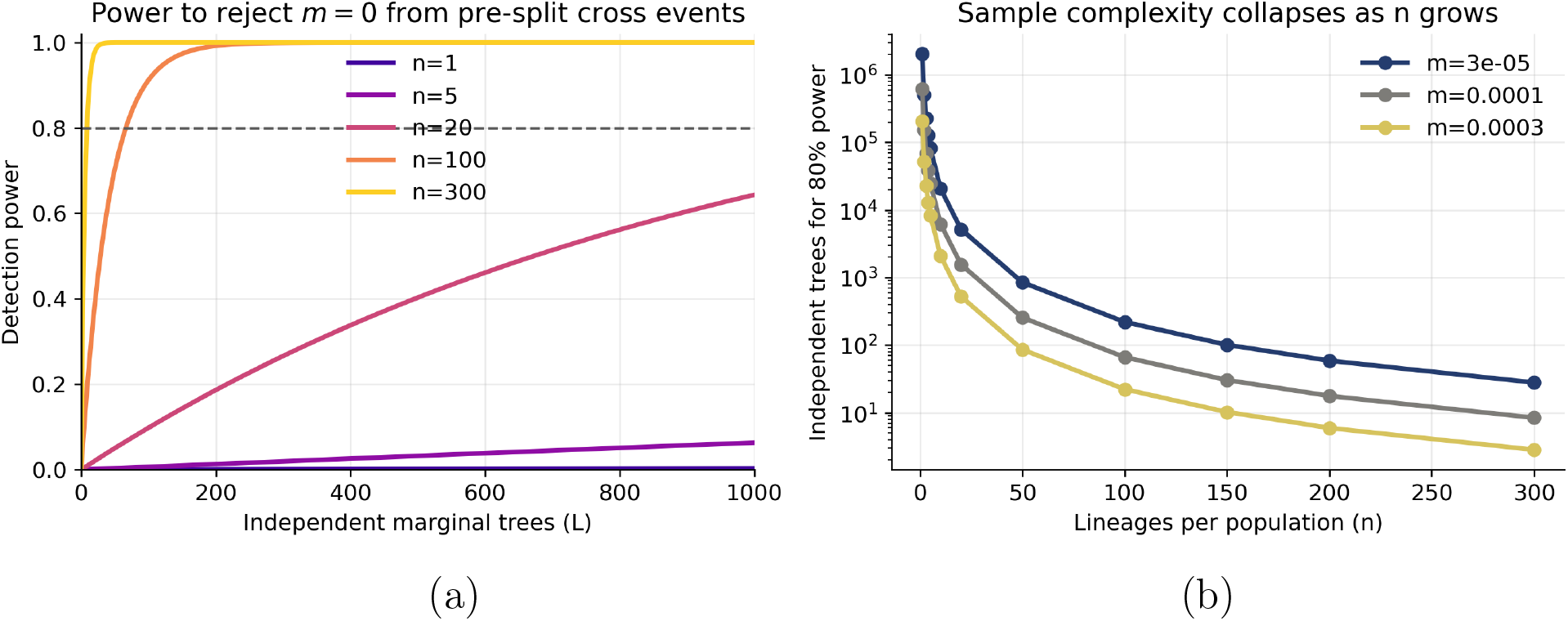
Power calculations for recent migration in IwM model with *T*_*s*_ = 30 generations. (a) Exact power 1 *−* (1 *− p*_*n*_)^*L*^ as a function of the number of independent marginal trees *L* for several balanced sample sizes. (b) The number of independent trees needed for 80% power as a function of *n*, shown for three migration rates.

### 3.4 Likelihood-based inference benchmarks

In these benchmarks, we assume that an ancestral recombination graph (ARG) is already available, either because it is observed directly in simulation or because it has been inferred from sequence data. We select *n* sampled *k*-tuples of haplotypes. Let *a*_*i*_ = (*a*_*i*1_, …, *a*_*ik*_) record the sampling demes for tuple *i*, and let *t*_*ij*_ be its first-coalescence time in marginal tree *j*. We use *T* sufficiently separated marginal trees and treat the resulting observations as independent. The resulting composite log-likelihood is

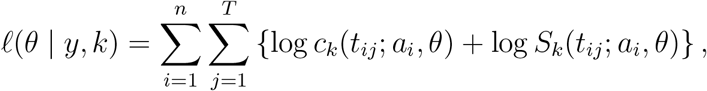

where *y* denotes the collection of first-coalescence times and sampling configurations. This use of summaries of coalescence times follows recent IICR-based inference work (Arredondo et al., 2021; DeHaas et al., 2025). We maximized the objective using SciPy’s `minimize` function with the `trust-constr` method (Virtanen et al., 2020); all parameter ranges are given in Table 1. We considered two benchmark settings: the `stdpopsim Africa_1T12` exponential growth model and a two-epoch extension of that growth history. We simulated observed ARGs with `msprime` and, for comparison, inferred ARGs from the same sequence data using `tsinfer` (Kelleher et al., 2019a) and `tsdate` (Pope et al., 2026). For the single-deme benchmarks based on `Africa_1T12`, we simulated 5 *×* 10^7^ base pairs for 100 haploid samples using the default model parameters and mutation rate 2.36 *×* 10^*−*8^. For each of 20 simulations, we drew 15 sets of evenly spaced marginal trees. Each set contained *n* = 500 randomly sampled *k*-tuples, with every tuple evaluated in each tree. We chose *T* = 400 for *k* = 2, using 125 kb spacing, and *T* = 100 for *k* = 50, using 500 kb spacing. The composite likelihood treats these observations as independent; in particular, it ignores dependence among tuples evaluated in the same tree. Each fit used 100 random initializations of the optimizer, after which we retained the run with the lowest negative log-likelihood; estimates from each simulation were then summarized by the median across the 15 tree sets. To construct inferred ARGs, we applied `tsinfer` followed by `tsdate` with default settings and mutation rate 2.36 *×* 10^*−*8^. The two-epoch growth model used the same inference pipeline but replaced the single recent growth phase by two growth epochs.

**Table 1:**
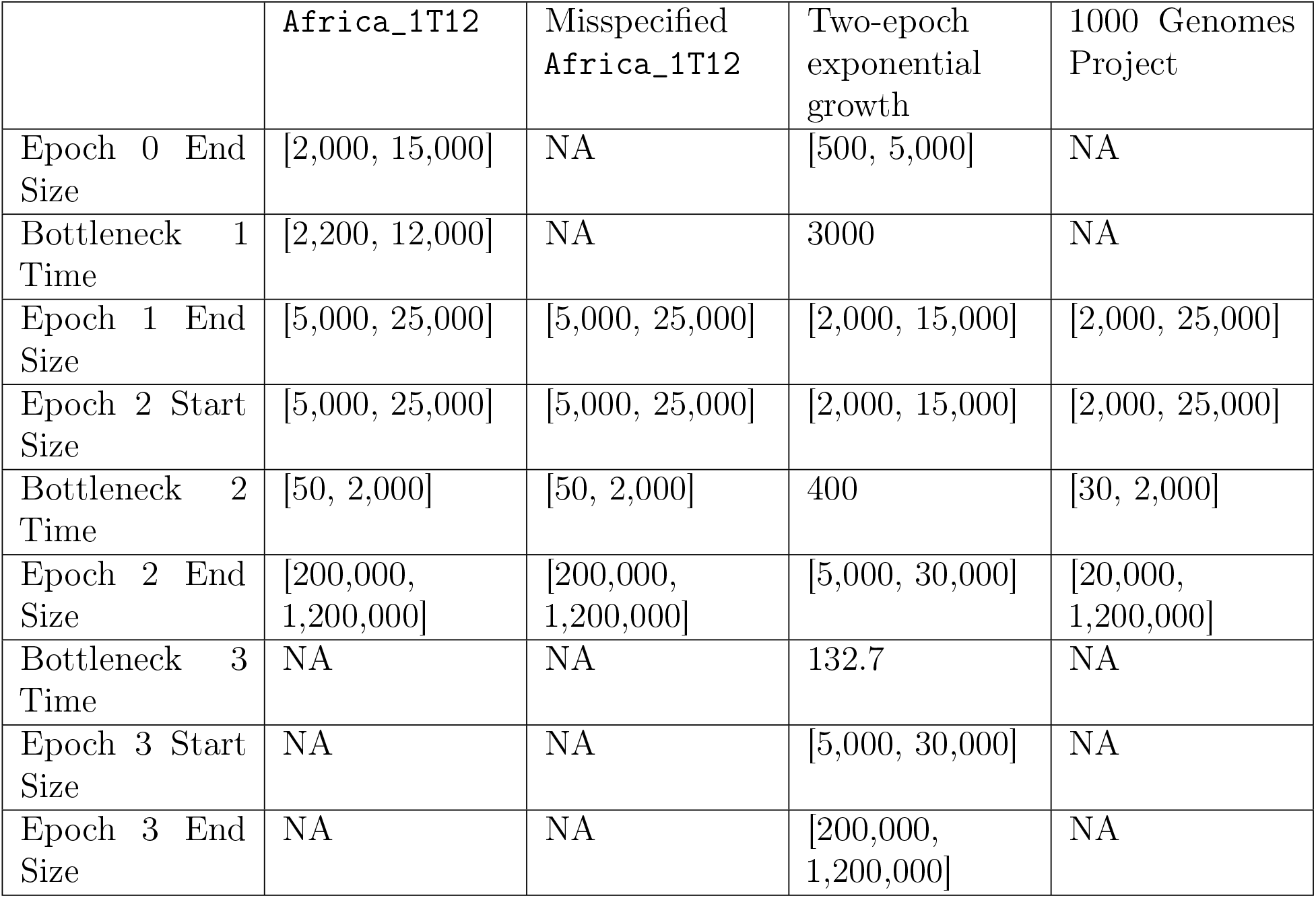
Parameter bounds and fixed values used for optimization. Bracketed entries are bounds, single values were fixed, and NA denotes parameters absent from the model. Wider bounds may require additional initializations because the objective is nonconvex.

#### 3.4.1 Single-epoch growth benchmark

Recent exponential growth concentrates first coalescences near the present, making present-day size difficult to learn from pairwise data alone. With the observed ARG, ICR_2_ recovers the older parameters accurately but has high error for present-day size (Figure 7). The absolute parameter estimates are presented in Appendix Figure 12.

**Figure 7:**
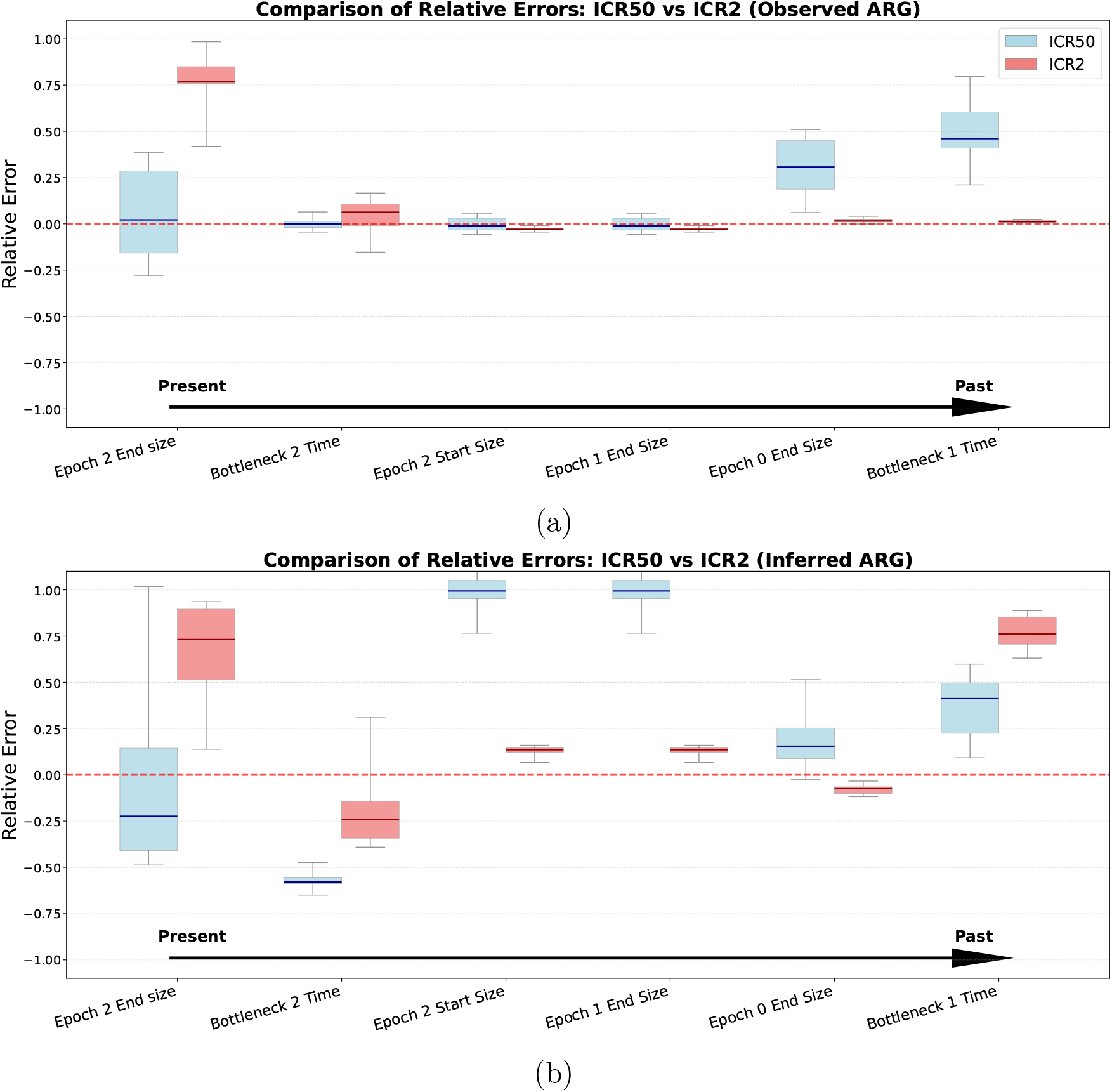
Relative error in inferred demographic parameters using (a) observed ARGs and (b) inferred ARGs. Parameters are ordered chronologically from the present to the past. With the observed ARG, ICR_2_ (red) is more accurate for older parameters, whereas ICR_50_ (blue) is more accurate for present-day size. With an inferred ARG, errors increase overall, but ICR_50_ remains more accurate than ICR_2_ for recent parameters.

Increasing the sample size to *k* = 50 raises the initial coalescence rate by a factor of 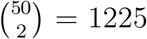, shifting information toward the recent bottleneck and present-day size at the expense of older parameters (Figure 7). We spaced sets of marginal trees by at least 125,000 base pairs for *k* = 2 and 500,000 base pairs for *k* = 50 to reduce linkage disequilibrium among first-coalescence times.

With inferred ARGs, ICR_50_ still estimates present-day size more accurately than ICR_2_ (Figure 7, lower). Figure 8 identifies the main error: inferred *T*_50_ values shift toward the present even though inferred *T*_2_ remains close to the truth. This distortion improves the fit to the latest epoch but biases bottleneck timing forward and degrades older estimates.

**Figure 8:**
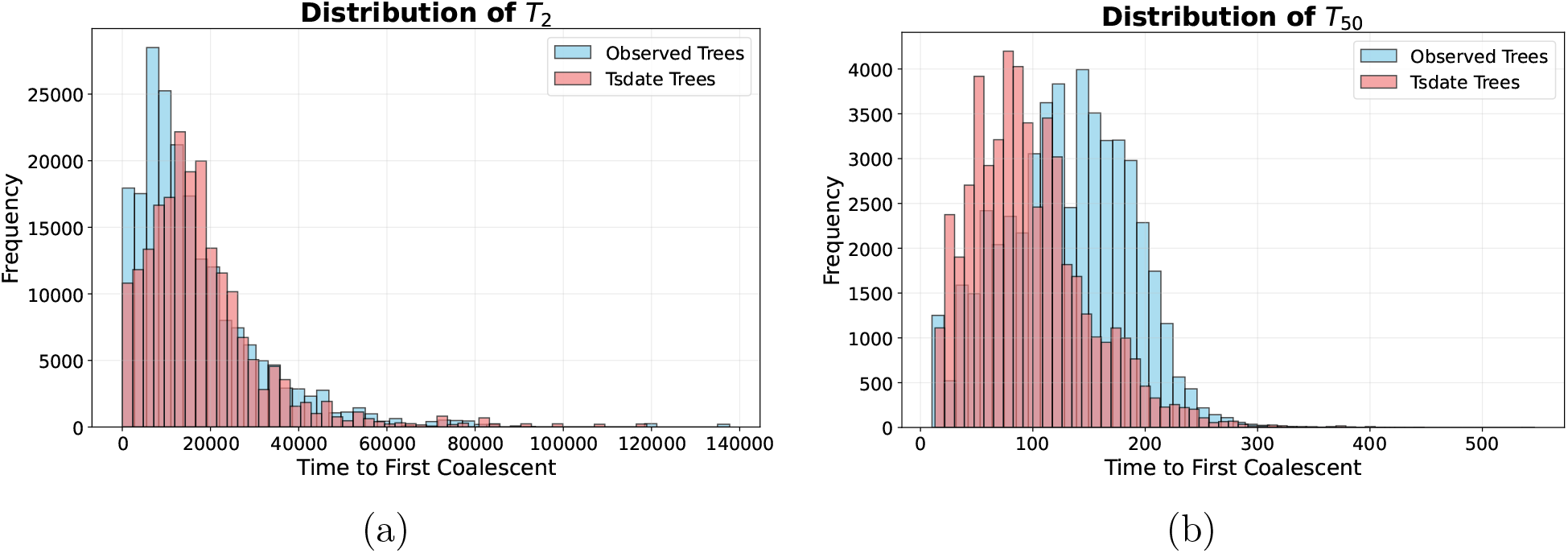
Observed and inferred distributions of *T*_2_ and *T*_50_ from one simulation. (a) The inferred *T*_2_ distribution is close to the truth, whereas (b) the inferred *T*_50_ distribution is shifted strongly toward the present.

Appendix Figure 17 repeats this comparison with `Relate`. In that example, both `Relate` and `tsinfer+tsdate` shift the *T*_50_ distribution toward the present, although the methods also differ in their recovery of *T*_2_.

#### 3.4.2 Inference under model misspecification

To test whether large-*k* inference of recent parameters tolerates a misspecified ancient history, we fitted a shortened version of the `Africa_1T12` model containing only presentday size and the recent bottleneck.

With the observed ARG, *k* = 50 recovers recent parameters despite the omitted deep history, whereas the pairwise fit does not (Figure 9). With inferred ARGs, bottleneck timing is biased but present-day size remains stable. These results should be interpreted in light of the shift in inferred *T*_50_ values shown in Figure 8.

**Figure 9:**
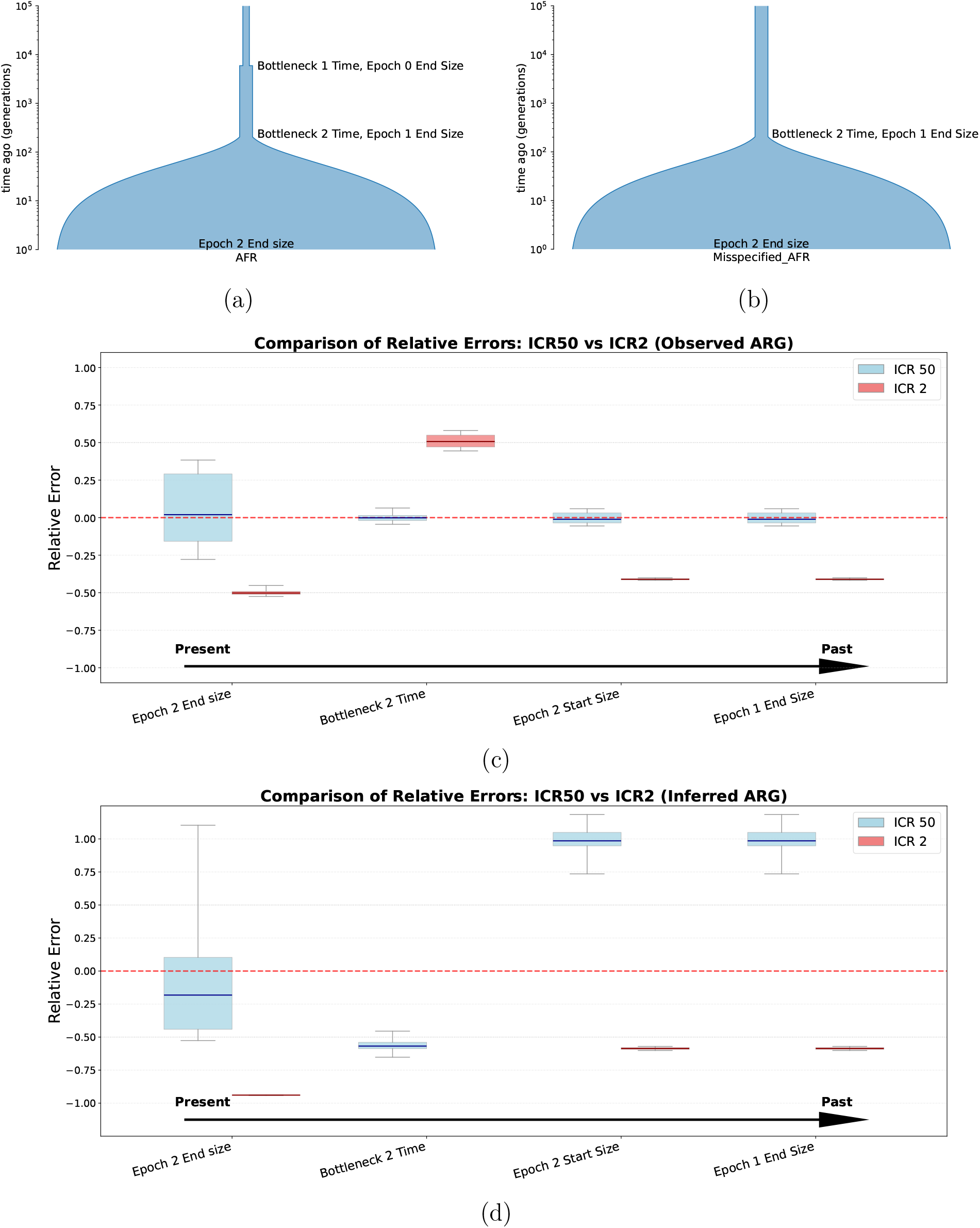
Inference under model misspecification. (a) Simulated African demographic model. (b) Fitted model, which omits ancient events. Fits to the observed (c) and inferred (d) ARGs show that ICR_50_ retains greater accuracy for recent parameters than ICR_2_. The ICR_2_ bottleneck estimate in (d) lies above the plotting range. Appendix Figure 13 gives the absolute estimates.

#### 3.4.3 Two-epoch exponential growth

We also fitted a two-epoch growth history based on the model of Gazave et al. (2014), fixing the event times and inferring only the size parameters. Estimates are most accurate for recent sizes, with greater error for the oldest size parameter (Figure 10).

**Figure 10:**
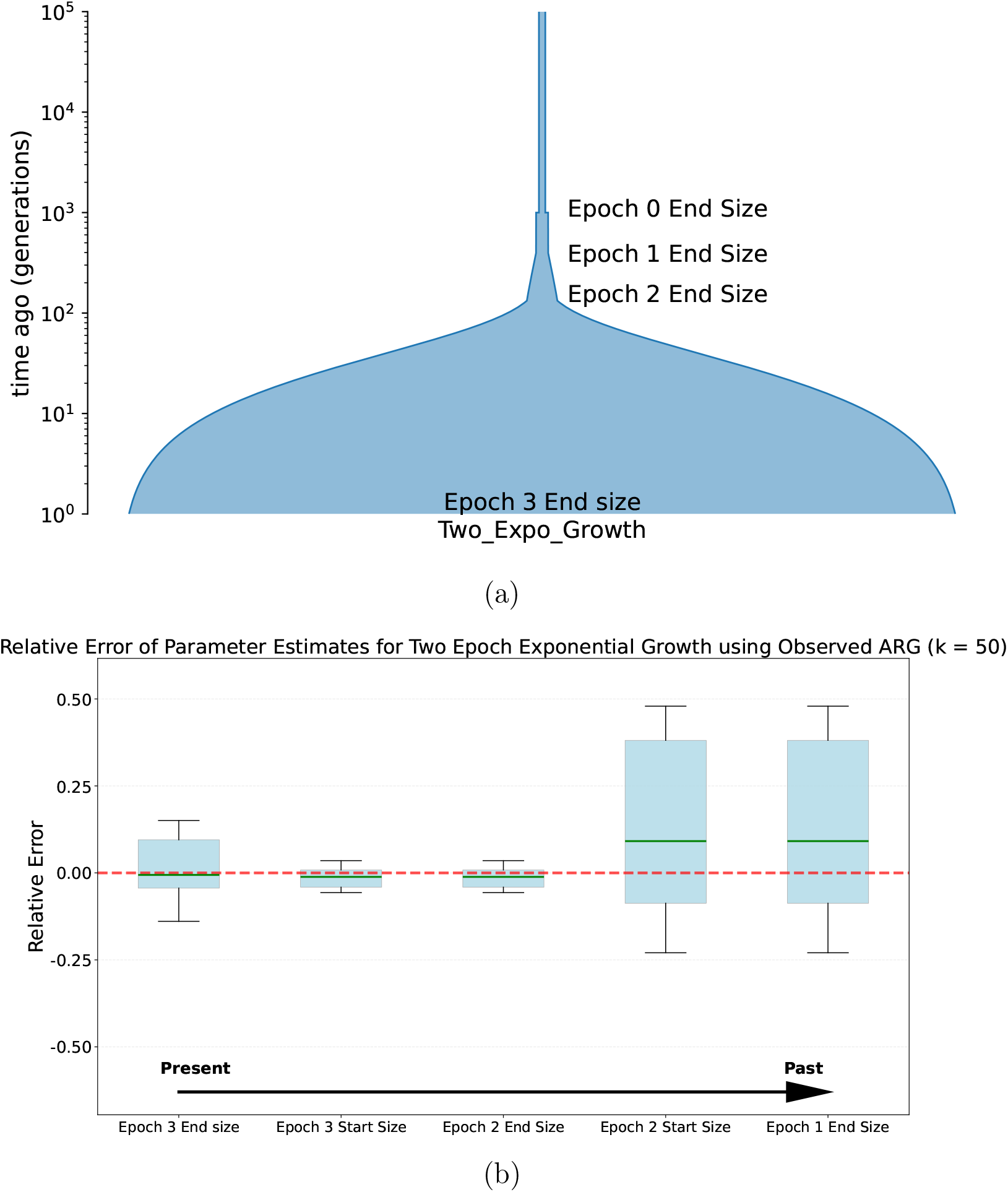
(a) the `Africa_1T12` model extended to two epochs of exponential growth. (b) Relative errors in estimates of population size. The ancient size is overestimated and exceeds the plotting range; Appendix Figure 14 gives the absolute estimates.

**Figure 11:**
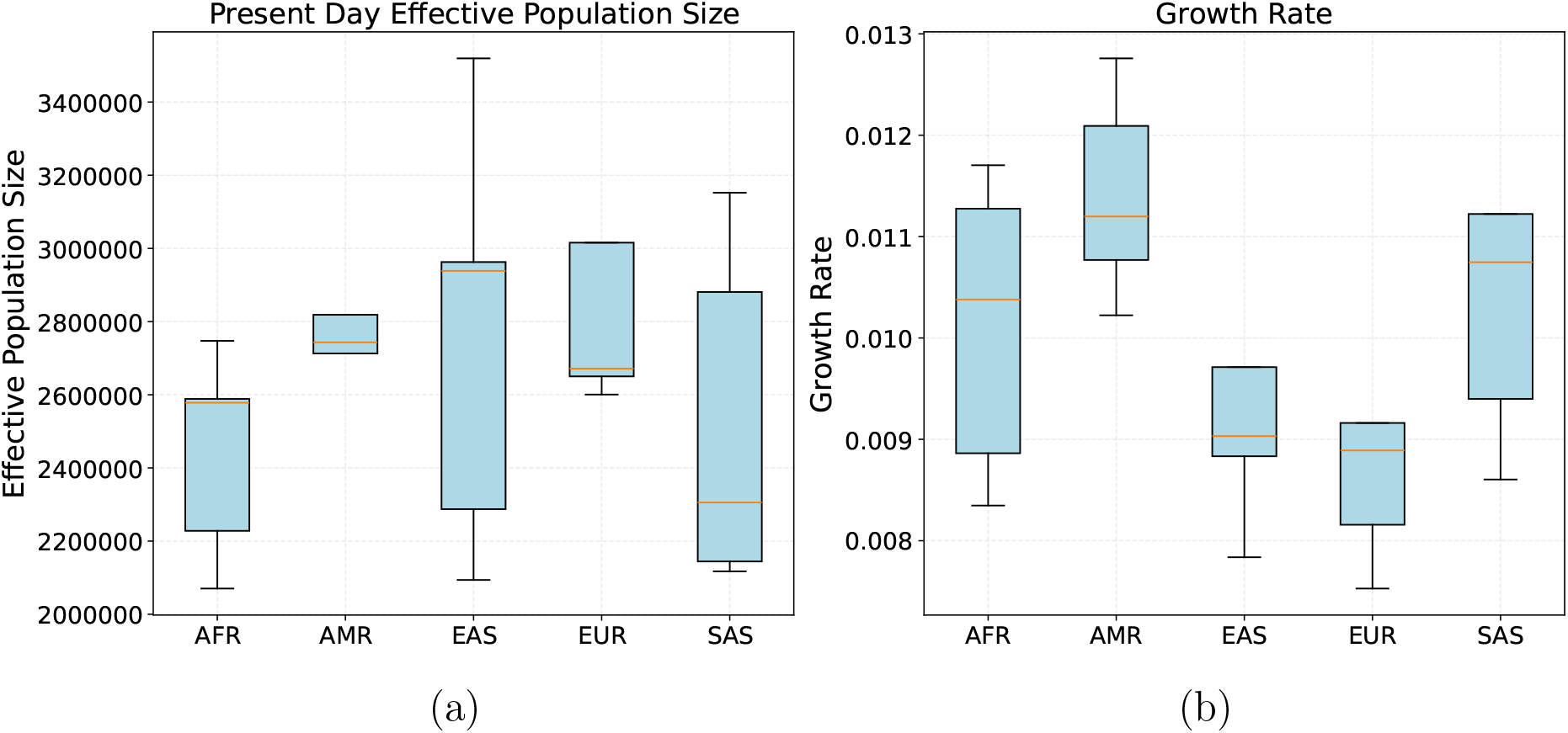
Inferred present-day effective population sizes (a) and exponential growth rates for five superpopulations. Each boxplot contains estimates from individual p arms of chromosomes 1 through 5.

Performance remains strongest in the recent part of the history. When event times are also allowed to vary, relative errors increase because more parameters must be estimated jointly. Appendix Figure 14(b) shows that performance also degrades on inferred ARGs, where `tsinfer` and `tsdate` shift *T*_50_ toward the present and thereby underestimate recent population sizes. This comparison should therefore be interpreted cautiously because the inferred *T*_50_ distribution is displaced before fitting.

### 3.5 Recent growth in 1000 Genomes populations

Estimates of recent human growth from the SFS vary across studies (Coventry et al., 2010; Gao and Keinan, 2016; Gazave et al., 2014; Nelson et al., 2012; Tennessen et al., 2012). Such estimates rely on rare allele frequencies and can depend on how ancient history is modeled. In our simulations, *k* = 50 first-coalescence times concentrate information in recent epochs, allowing the ancient part of the model to be simplified, although accuracy still depends on the inferred ARG.

We fitted the model of recent growth in Figure 9(b) to tree sequences estimated by Wohns et al. (2022), using *k* = 50. Following Gazave et al. (2014), we restricted the model to recent parameters. We included populations with at least 100 haploid samples and fitted each population separately; for each chromosome arm, we summarized each superpopulation by the median of its population estimates. For the European (EUR) samples, we estimate a growth rate of about 0.9% per generation and a present-day effective population size of about 2.7 million. The growth rate is lower than the estimate in Gazave et al. (2014) and closer to those reported by Nelson et al. (2012). Across the five superpopulations, the present-day effective sizes are estimated to be on the order of millions, and the fitted recent growth rates range from about 0.9% to 1.1% per generation. The onset of growth is placed 250–350 generations ago, or about 6,250–8,750 years ago assuming a 25-year generation time. Appendix Figure 15 gives the corresponding estimates. Results from the q arms of EUR chromosomes are similar (Appendix Figure 16).

Finally, these estimates may be biased because they depend on inferred ARGs. In our simulations, `tsinfer` and `tsdate` shifted inferred *T*_50_ values toward the present. This led to an overestimate of the/ population size before growth and suggested that the duration of growth and present-day size may be underestimated (Figure 7). Similar errors in the 1000 Genomes tree sequences would bias the estimates in Figure 11 in the same directions. As a sensitivity check, replacing our inferred size before growth with the estimate from Gazave et al. (2014), while retaining our present-day size and the inferred time at which growth began, gives a growth rate of 1.82%.

## 4 Discussion

In this paper we introduced `demestats`, a method for computing first-coalescence and cross-coalescence statistics for arbitrary sample sizes and sampling configurations. Exact calculations are supplemented by a mean-field approximation for large samples. These statistics quantify parameter sensitivity under a given sampling design and improve resolution of recent size changes and migration.

Our simulation results show that pairwise summaries are often most informative about older parts of the demography, whereas first-coalescence statistics from larger samples concentrate information near the present. In our benchmarks, this improves inference for recent growth and recent migration, and it also makes inference less sensitive to misspecification of ancient events when the parameters of interest are recent. In the 1000 Genomes analysis, the same approach gives present-day effective sizes and growth rates that are broadly consistent with earlier SFS-based studies.

The main limitation of our analyses is their dependence on the quality of the input tree sequence. In our simulations, the ARG inference method we used (`tsinfer`+`tsdate`) recovered the distribution of *T*_2_ more closely than that of *T*_50_, which was shifted toward the present. A comparison with `Relate` showed the same recentward shift for *T*_50_ in this benchmark (Appendix Figure 17). Better ARG inference, or corrections for this distortion, would directly improve inference with `demestats`. Another open problem is how best to combine small-*k* and large-*k* likelihoods so that both ancient and recent demographic parameters can be estimated accurately in the same analysis.

## Data and Code Availability

The inferred tree sequences used in this paper come from the unified genealogy of modern and ancient genomes (Wohns et al., 2022) and are publicly available through the accompanying 1000 Genomes tree sequence release (Kelleher et al., 2019b). Before analyzing them, we applied the latest available version of `tsdate` (v0.2.6; Pope et al., 2026) for improved time calibration.

`demestats` is implemented in Python with documentation and tutorials available at https://demestats.readthedocs.org and the code is publicly available at https://github.com/jthlab/demestats.

## Acknowledgments

This research was supported in part by the National Institute of General Medical Sciences of the NIH under award number R35GM151145. The content is solely the responsibility of the authors and does not necessarily represent the official views of the NIH.

## A Exact ICR transition matrices

Consider one block with demes indexed by *i* = 1, …, *d* and *m* lineages whose first coalescence has not yet occurred. Between demographic events, the exact ICR calculation has the form

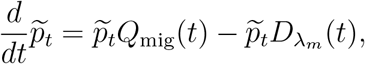

with the same row-vector convention as in (1). Before allocating an array, the implementation compares simple estimates of the storage required by two equivalent encodings of the state and chooses the lower one.

### Occupancy representation

The occupancy state space is

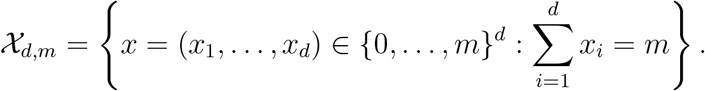

Here *x*_*i*_ is the number of lineages in deme *i*. Let *e*_*i*_ denote the *i*th standard basis vector, and let *M*_*ij*_(*t*) be the backward-time lineage migration rate from deme *i* to deme *j*. Then

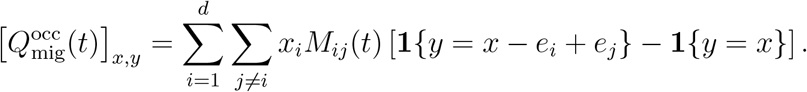

Thus, for each ordered pair (*i, j*) with *i* ≠ *j*, one lineage moves from deme *i* to deme *j* at rate *x*_*i*_*M*_*ij*_(*t*), and the diagonal entry is the negative sum of outgoing migration rates. The coalescence rate in state *x* is

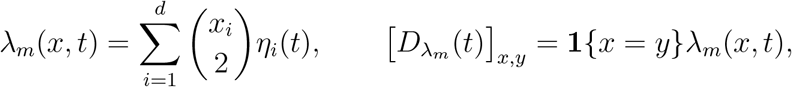

where *η*_*i*_(*t*) = 1*/{*2*N*_*i*_(*t*)*}*.

#### Labeled-lineage representation

The labeled-lineage formulation keeps the original lineage labels and uses the extra symbol *⊥* for “outside the current block.” If the full sample has size *k* and the current block is *B* = *{*1, …, *d}*, the implemented state space is

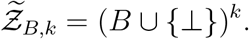

Here *z*_*l*_ = *i ∈ B* means that lineage *l* is represented in this block and currently lies in deme *i*, while *z*_*l*_ = *⊥* means that lineage *l* is not represented in this block. If the block currently carries *m* lineages, then restricting to states with exactly *m* non-*⊥* coordinates and dropping the *⊥* entries recovers the equivalent *B*^*m*^ description used in Section 2. Keeping all *k* label positions allows two block distributions to be multiplied when the blocks join: each lineage is present in one incoming block and marked *⊥* in the other.

For disjoint blocks *B*_1_ and *B*_2_, let 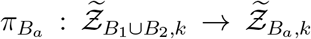 send every coordinate outside *B*_*a*_ to *⊥*. If donor deme *a* is merged into recipient deme *b* within one block, let *µ*_*a→b*_(*z*) denote the state obtained from *z* by replacing every coordinate equal to *a* by *b*. For `Split2`, let *B*_donor_ and *B*_recipient_ denote the donor and recipient blocks. Let *T*_*B*_ be the time of the first coalescence among the *m* lineages represented in the current block. In the formulas below, *p* and *p*^*′*^ are the normalized distributions of lineage locations conditional on *T*_*B*_ not having occurred, immediately younger and older than the demographic event, respectively. The symbols *p*^(1)^ and *p*^(2)^ denote the corresponding distributions from two incoming blocks. Survival probabilities are stored separately. In the `Pulse` and `Admix` formulas below, *u* runs over all valid incoming labeled-lineage states. Events are updated as follows:

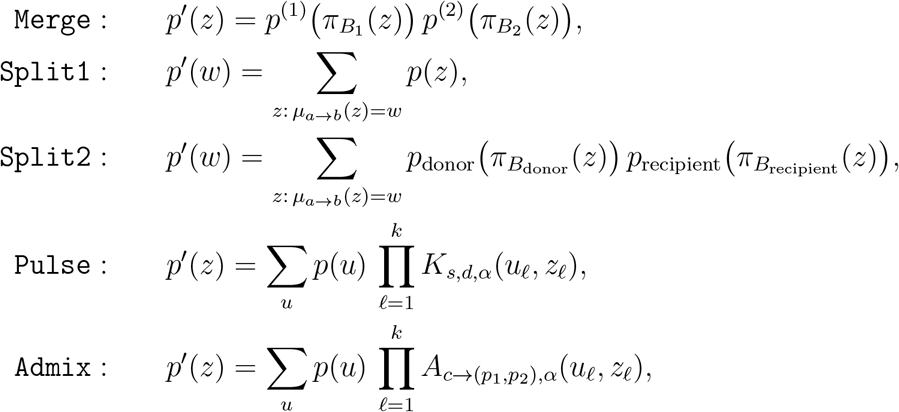

where a pulse with source *s*, destination *d*, and proportion *α* uses the following transition probabilities:

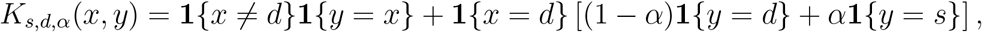

and an admixture event with child *c*, parents *p*_1_, *p*_2_, and proportion *α* uses

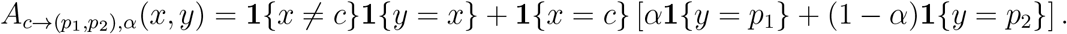

Thus, pulse and admixture updates are applied independently to each labeled lineage, whereas `Merge`, `Split1`, and `Split2` are deterministic relabelings combined with multiplication across independent incoming blocks.

Between demographic events, define 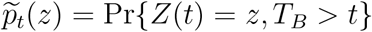. This distribution obeys

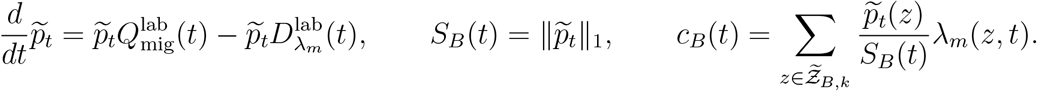

If *n*_*i*_(*z*) = #*{l* : *z*_*l*_ = *i}* for *i ∈ B*, then

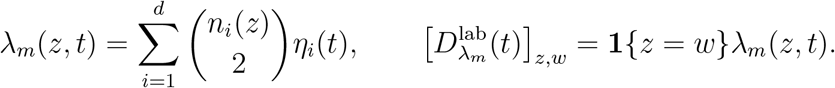

When the block contains all *k* lineages, *T*_*B*_ = *T*_*k*_, *S*_*B*_ = *S*_*k*_, and *c*_*B*_ = *c*_*k*_. Before then, survival probabilities from independent blocks are multiplied when the blocks join. If M(*t*) is the migration rate matrix for a single lineage on *B ∪ {⊥}*, with entries

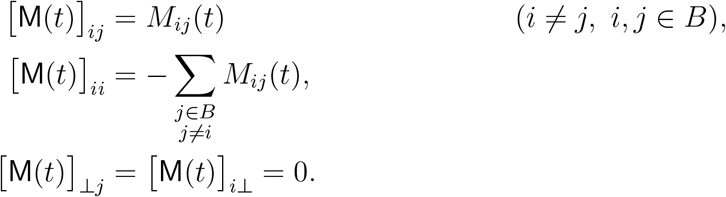

then

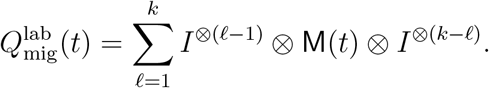

Here *I* is the identity matrix and *⊗* denotes the Kronecker product. Each term lets lineage *l* migrate while leaving the locations of the other lineages unchanged; their sum lets all labeled lineages migrate independently.

On a constant-rate interval, either representation uses the matrix exponential of the full transition-rate matrix. On an interval with time-varying migration or population sizes, the implementation solves the corresponding differential equation directly.

## B Exact CCR transition matrices

On a constant-rate interval with demes indexed by *i* = 1, …, *d* and total sample size *k*, the exact CCR calculation evolves the probabilities 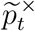 defined in the main text on the count state space

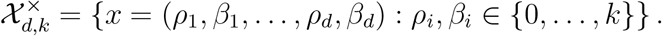

Here *ρ*_*i*_ and *β*_*i*_ are the integer numbers of red and blue lineages in deme *i*. The direct implementation allocates an array with (*k* + 1)^2*d*^ entries, although only a reachable subset carries nonzero mass. Let 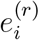 and 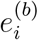 denote the unit vectors on the red and blue coordinates of deme *i*, let *M*_*ij*_(*t*) be the backward-time lineage migration rate from deme *i* to deme *j*, and let *η*_*i*_(*t*) = 1*/* {2*N*_*i*_(*t*)}. With the row-vector convention used in (2), the transition-rate matrix has the following entries.

The migration matrix for red lineages moves one lineage from deme *i* to deme *j*:

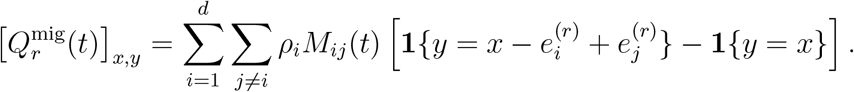

Equivalently, for each ordered pair (*i, j*) with *i* ≠ *j*, the only nonzero off-diagonal transition is

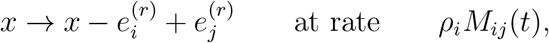

and the diagonal entry is the negative sum of outgoing migration rates for red lineages. The blue migration matrix is defined analogously, with *β*_*i*_ in place of *ρ*_*i*_ and 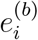 in place of 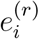.

The red-red coalescence matrix records coalescence of two red lineages into one surviving red lineage in the same deme:

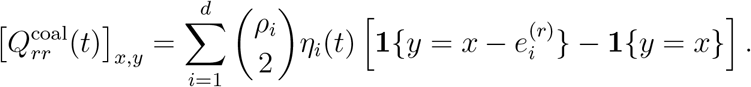

Thus, if *ρ*_*i*_ *≥* 2, the state jumps from *x* to 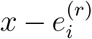 at rate 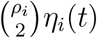, reflecting that two red lineages are replaced by one red ancestor. The blue-blue coalescence matrix is defined analogously, with *β*_*i*_ in place of *ρ*_*i*_ and 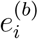 in place of 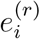.

Finally, a red-blue coalescence ends the event being followed and therefore removes probability at rate

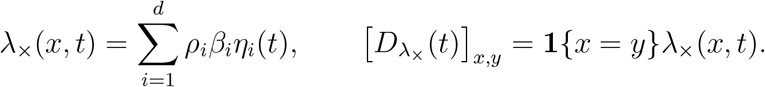

The matrix governing the survival-weighted probabilities on a constant interval is therefore

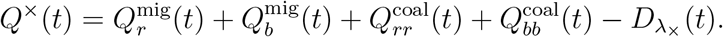

## C Additional figures

**Figure 12:**
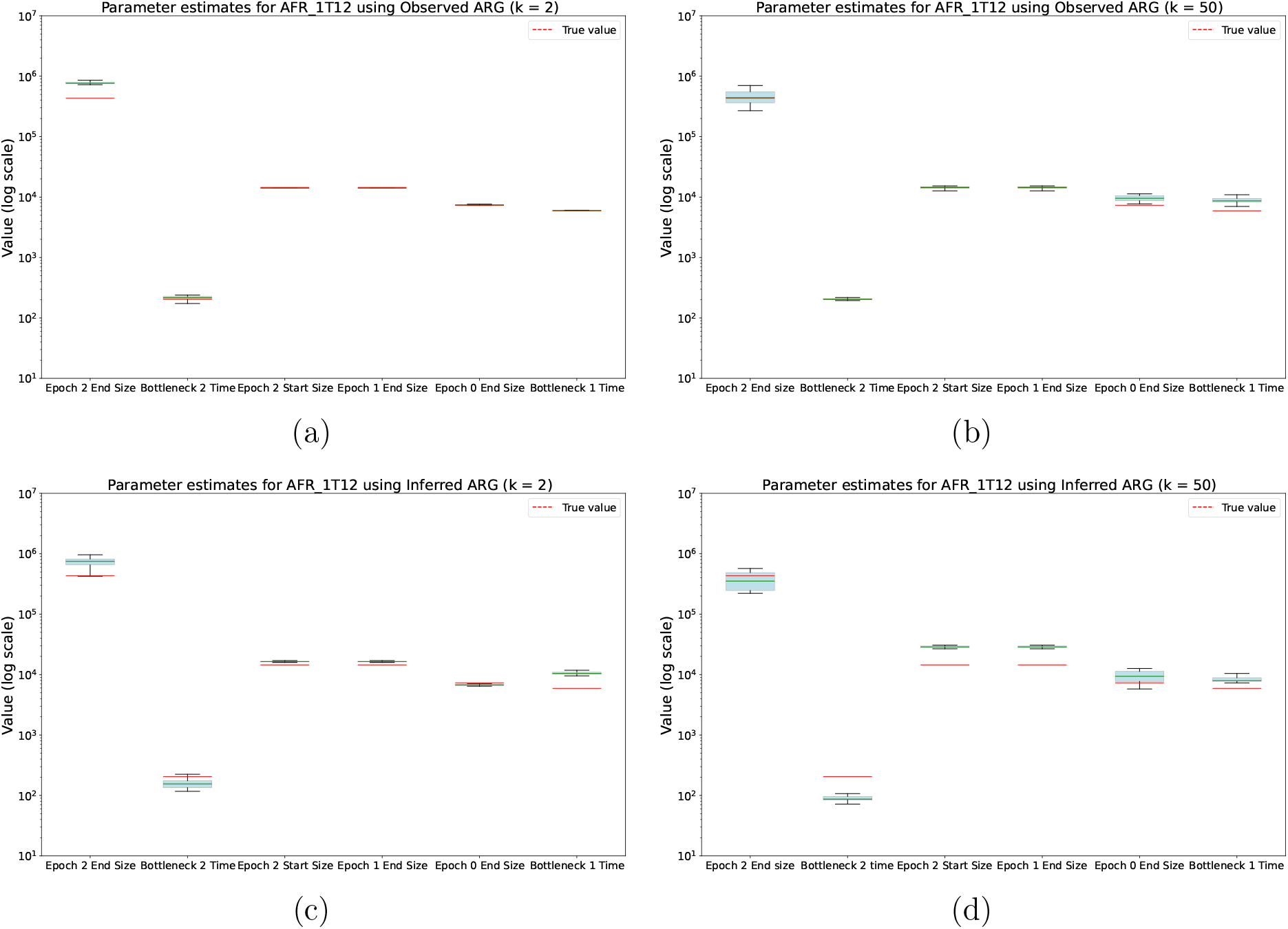
Parameter estimates for the `Africa_1T12` model from observed ARGs (top) and inferred ARGs (bottom), with *k* = 2 on the left and *k* = 50 on the right.

**Figure 13:**
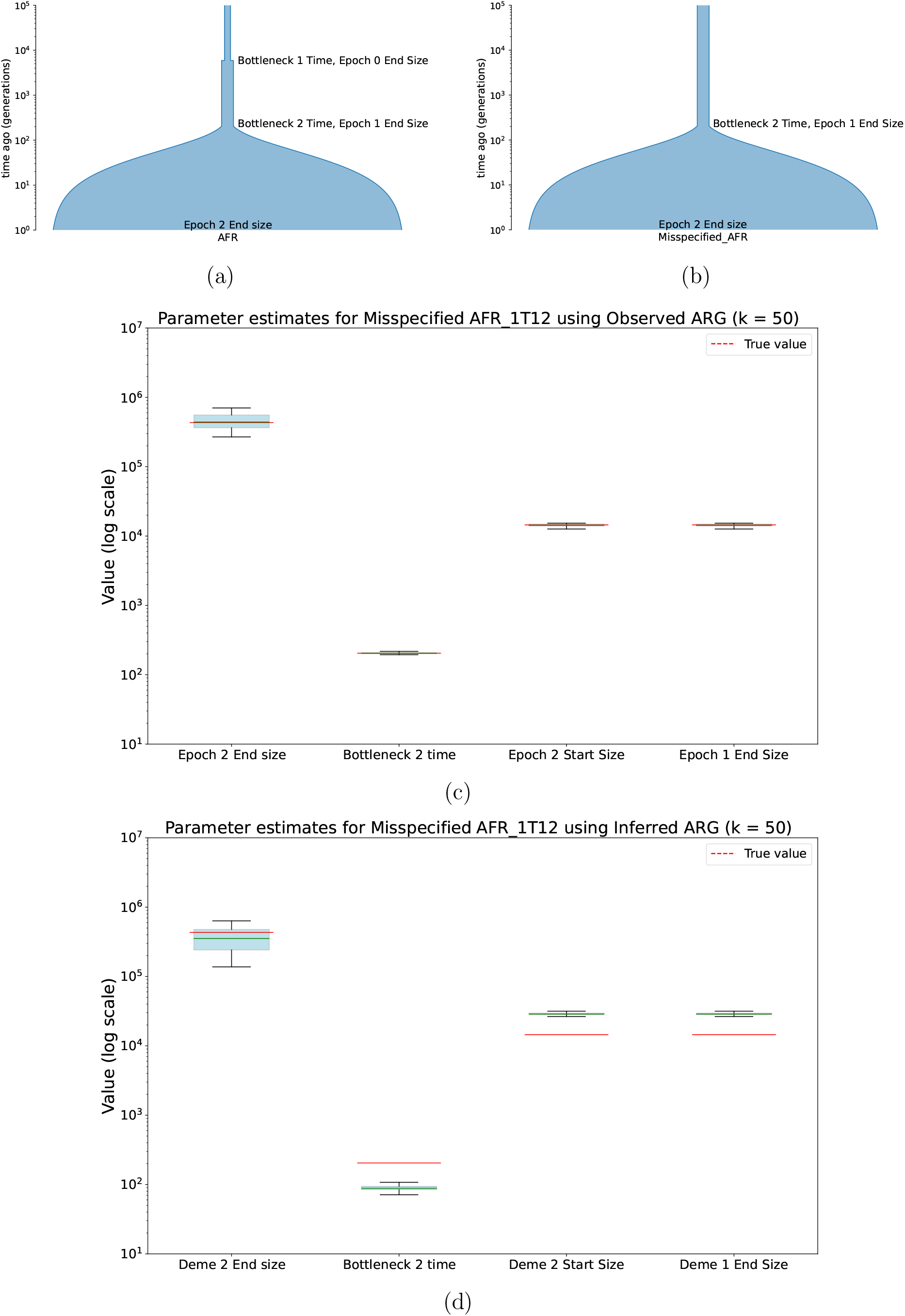
Inference under model misspecification. The original `Africa_1T12` model (a) and misspecified model (b) are followed by fits to observed (c) and inferred (d) ARGs.

**Figure 14:**
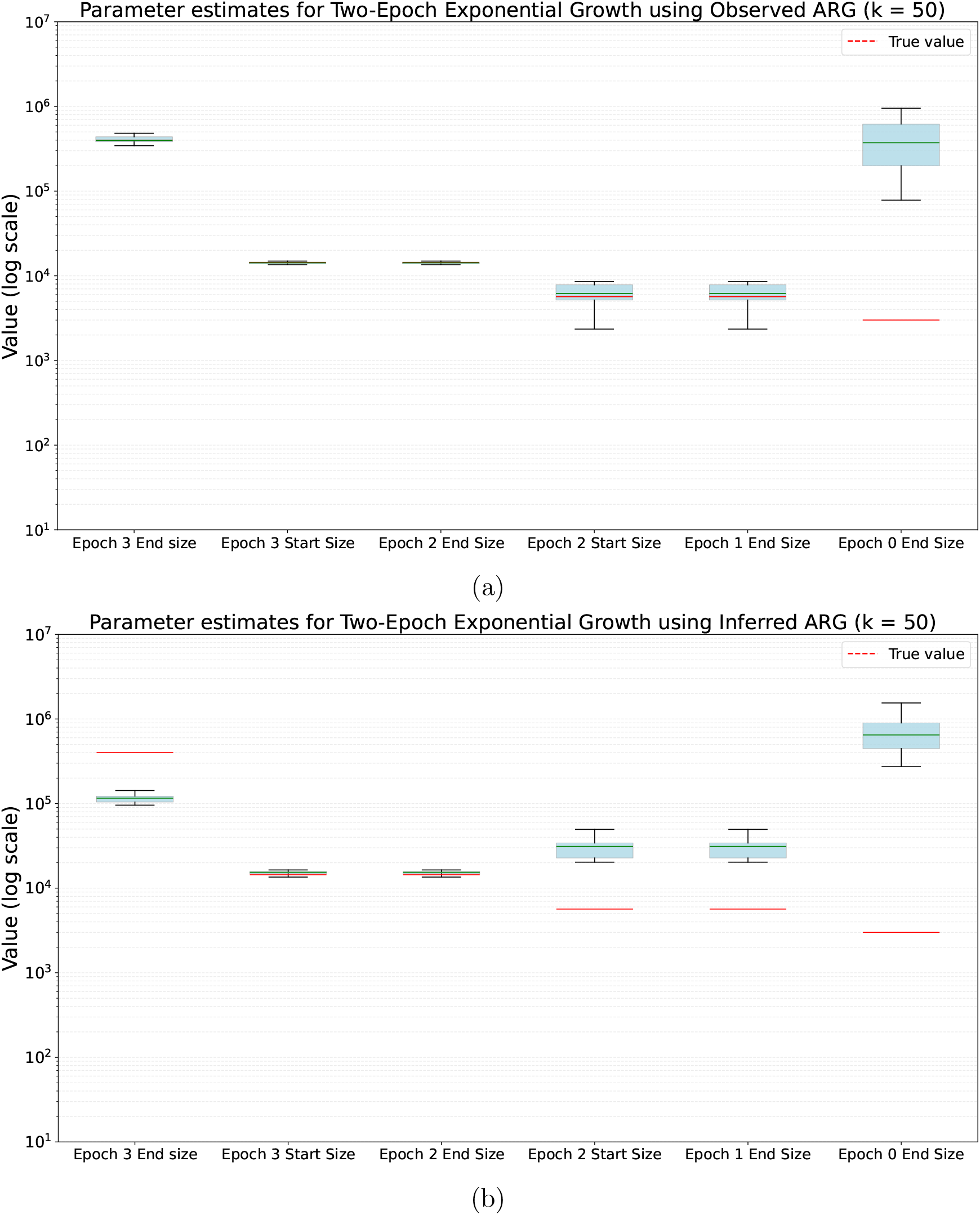
Parameter estimates for the two-epoch exponential growth model using ICR_50_, from observed (a) and inferred (b) ARGs. Errors are larger for the inferred ARG, consistent with the recentward shift in inferred *T*_50_ values.

**Figure 15:**
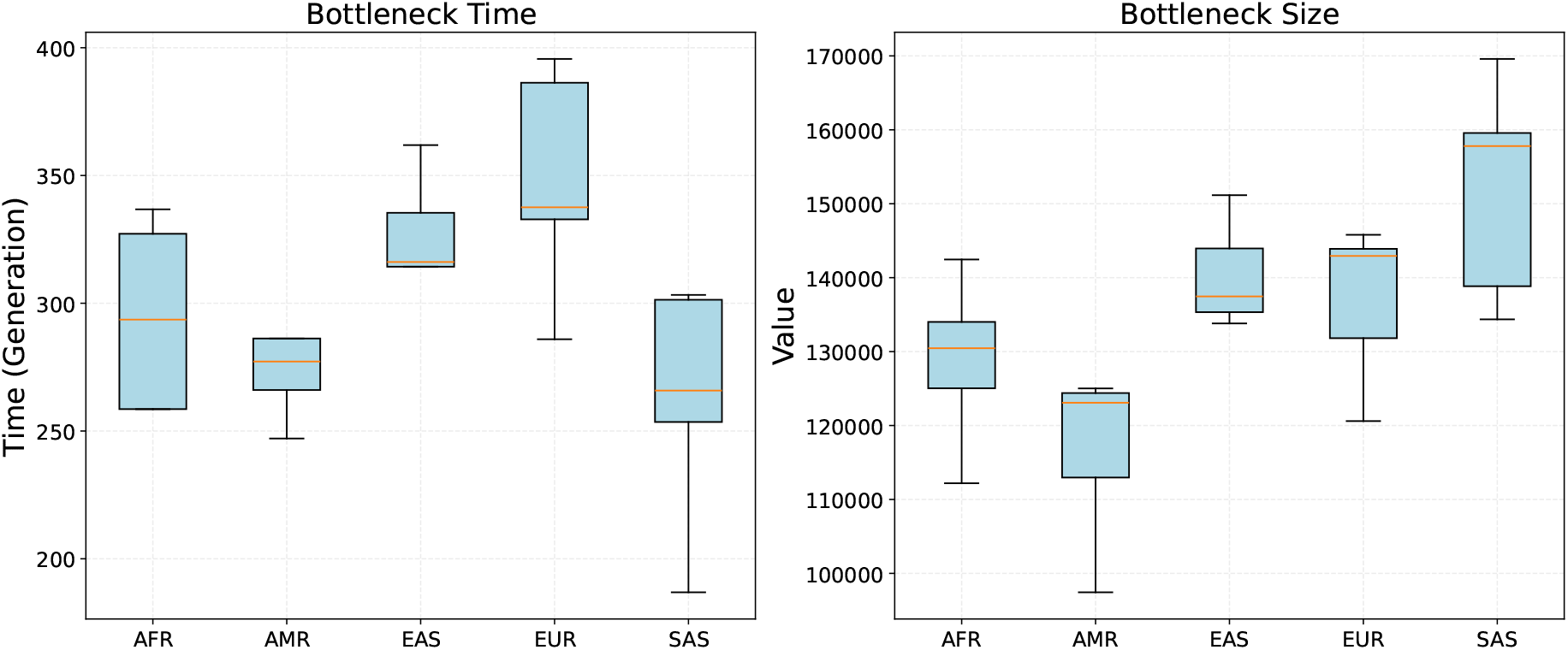
Inferred bottleneck times and sizes for five superpopulations. Each boxplot contains estimates from the p arms of chromosomes 1–5.

**Figure 16:**
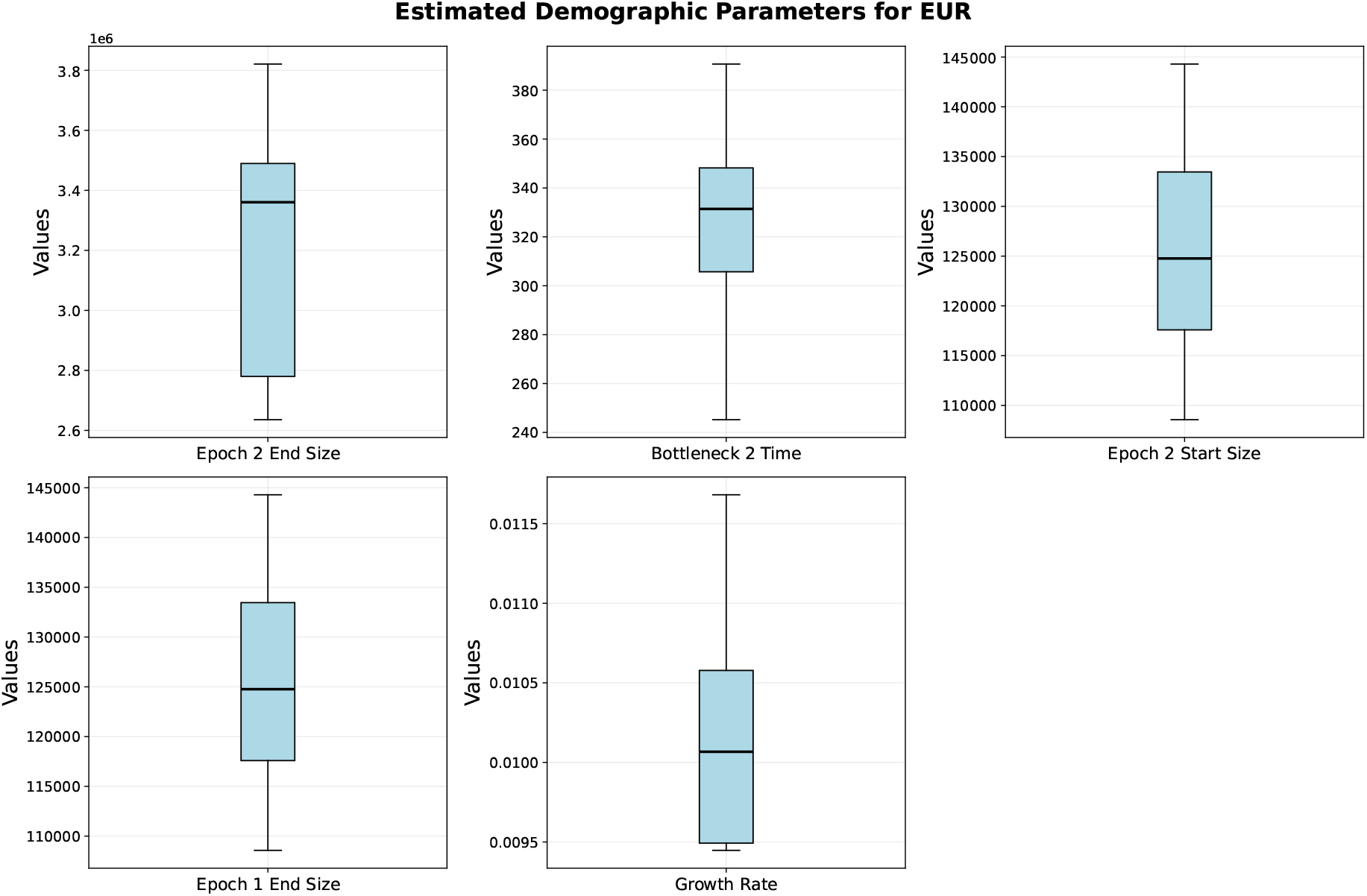
EUR demographic parameter estimates from the q arms of chromosomes 1–10. The results are similar to the p-arm estimates in Figure 11.

**Figure 17:**
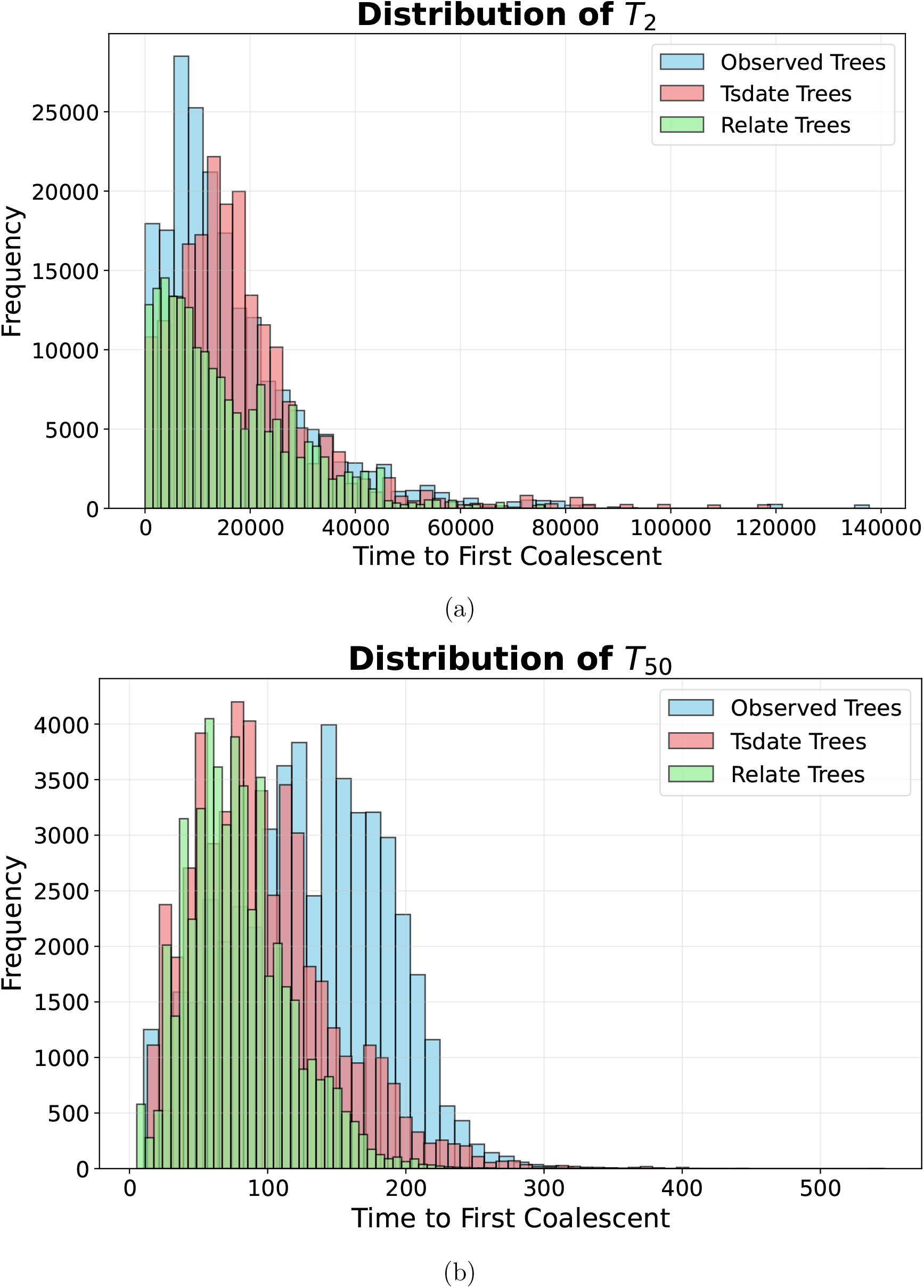
Observed, `tsdate`, and `Relate` distributions of (a) *T*_2_ and (b) *T*_50_ from one simulation. Both inferred *T*_50_ distributions are shifted toward the present, although the methods differ in their recovery of *T*_2_.

## Notes

### Competing Interest Statement

The authors have declared no competing interest.

### Summary of Updates

Reworked the mean-field formulation to derive coalescence rates and lineage-count dynamics from an explicit assumption of independent lineage states conditional on survival, accounting for lineage-count variance and survival conditioning. Clarified the introduction and notation, added figure-panel labels, improved captions and simulation-parameter descriptions, and corrected minor errors.

